# Microbe tree metabolite interactions in the soil - phyllosphere continuum of poplar tree: when microbes rewire poplar root exudate and metabolome

**DOI:** 10.1101/2024.03.04.583280

**Authors:** F. Fracchia, F. Guinet, N.L. Engle, T.J. Tschaplinski, C. Veneault-Fourrey, A. Deveau

## Abstract

- Trees are associated with a broad range of microorganisms colonising the diverse tissues of their host. However, the early dynamics of the assembly of the microbiota from the root to shoot axis and how it is linked to root exudates and metabolite contents of tissues remain unclear.
- Here, we characterized how fungal and bacterial communities are altering root exudates as well as root and shoot metabolomes in parallel with their establishment in poplar cuttings (*Populus tremula x tremuloides* clone T89) over 30 days of growth. Sterile poplar cuttings were planted in natural or gamma-irradiated soils. Bulk and rhizospheric soils, root and shoot tissues were collected from day 1 to day 30 to track the dynamic changes of fungal and bacterial communities in the different habitats by DNA metabarcoding. Root exudates and root and shoot metabolites were analysed in parallel by gas chromatography-mass spectrometry.
- Our study reveals that microbial colonization triggered rapid and substantial alterations in both the composition and quantity of root exudates, with over 70 metabolites exclusively identified in remarkably high abundances in the absence of microorganisms. Noteworthy among these were lipid-related metabolites and defence compounds. The microbial colonization of both roots and shoots exhibited a similar dynamic response, initially involving saprophytic microorganisms and later transitioning to endophytes and symbionts. Key constituents of the shoot microbiota were also discernible at earlier time points in the rhizosphere and roots, indicating that the soil constituted a primary source for shoot microbiota.
- Furthermore, the microbial colonization of belowground and aerial compartments induced a reconfiguration of plant metabolism. Specifically, microbial colonization predominantly instigated alterations in primary metabolism in roots, while in shoots, it primarily influenced defence metabolism. This highlighted the profound impact of microbial interactions on metabolic pathways of plants, shedding light on the intricate interplay between plants and their associated microbial communities.

## INTRODUCTION

The plant-associated microbiota is considered as the second genome of the host plant. It comprises a diverse and complex range of microorganisms, including bacteria, archaea, fungi, oomycetes and viruses (Trivedi et al., 2020). Members of these microbial communities can either be detrimental, such as pathogens, or on the contrary, favourable to their host (Mendes et al., 2013). Thus, they play essential roles in plant life traits. They promote nutrient acquisition, plant growth (Compant et al., 2010; Richardson et al., 2009; Rodriguez et al., 2006; White et al., 2019), and resistance to biotic and abiotic stresses (Haney et al., 2015; Hassani et al., 2018). The microbiota activities can be seen as the extended phenotype of plants (Brown et al., 2020). Plants provide a multitude of habitats for the development and proliferation of microbial communities. Regardless of whether the plant is an annual or a perennial species, microbial community composition varies significantly between the bulk soil, rhizosphere, root endosphere and phyllosphere, indicating that the plant compartment is a major selective force for the assembly of the microbiota (Beckers et al., 2017; Bodenhausen et al., 2013; Brown et al., 2020; Daghino et al., 2022; Swift et al., 2021; Trivedi et al., 2020; Wallace et al., 2018; Wei et al., 2021).

These microbial communities are dynamic in time and space and their assembly is regulated by both biotic (e.g., host genotype, microbe-microbe interactions) and abiotic factors (e.g., soil origin, climate, seasonal variation). Soil provides the main reservoir for root microbial communities, while both vertical (via seeds) and horizontal transmission (via soil, air, insects and/or other plants) are sources for phyllosphere microbial colonisation (Vorholt, 2012). Nevertheless, the relative roles of soil and air pathways for phyllosphere colonisation are not yet clearly established with contradictory results. Some evidence suggests that phyllosphere microorganisms are sourced from the soil (Xiong et al., 2021), while other studies observe the air as the main reservoir (Maignien et al., 2014), or a dual influence (Dove et al., 2021). The rhizosphere is the first compartment where the host genotype starts to influence the microbiota composition through rhizodeposits (Reinhold-Hurek et al., 2015; Sasse et al., 2018). This selection of microbial communities between soil and rhizosphere has been largely documented and results in a decrease of diversity (Gottel et al., 2011; Lundberg et al., 2012). In the host endosphere, plant-microbe and microbe-microbe interactions are the main factors driving the assembly of the microbiota (Lareen et al., 2016), where its diversity decreases from belowground to aboveground compartments (Trivedi et al., 2020).

Although trees are long-lived perennials whose microbiota evolves over the course of their lives and the stage of forest cover (Xie et al., 2023), the initial assembly of the microbiota is thought to influence plant health and physiology (De Zelicourt et al., 2013; Dove et al., 2021; Mangeot-Peter et al., 2020). Previous studies have demonstrated that the microbiota of belowground and aboveground tissues of poplars (*Populus* sp.) change drastically over the first months of growth in natural soil (Dove et al., 2021; Mangeot-Peter et al., 2020), and that both selective and stochastic factors operate in the structuring of the poplar root microbiota (Dove et al., 2021). On a finer scale, we have previously shown that naive poplar roots are colonised within a few days and that several waves of fungi and bacteria follow one another over the first 50 days, with saprotrophs being slowly replaced by endophytes and symbionts (Fracchia et al., 2021). While the establishment of a molecular dialog between the root cells and the mycorrhizal and endophytic fungi likely explain the delayed colonisation by these fungi, other mechanisms presumably drive this colonisation. For example, root exudates are expected to play a key role in the chemoattraction of rhizospheric microbes and their habitat structuring (Chai and Schachtman, 2022; Haichar et al., 2008; Seitz et al., 2022; Zhalnina et al., 2018). Conversely, rhizospheric microbes can systemically modulate the composition of root exudates (Korenblum et al., 2020). The composition of root exudates also depends on the plant species, genotype, developmental stage, and environmental conditions, but all root exudates contain broadly the same classes of compounds derived from primary and secondary metabolisms: sugars, organic acids, amino acids, lipids, proteins, terpenes, phenolics, flavonoids (Badri and Vivanco, 2009). Most of the studies report root exudate composition using sterile hydroponic systems, followed by the characterization of the role of one type of metabolites in the interaction with microbiota. Furthermore, most studies have been conducted on herbaceous plants or shrubs, whereas similar studies with trees are limited to a few species and involve very few forest trees (Vives-Peris et al., 2020). To our knowledge, only one study has attempted to characterize poplar root exudates, and it focused more on rhizospheric soil metabolomes rather than on actual exudates (Li et al., 2022). Even less is known regarding feedback effects of rhizospheric microbiota on tree exudation. The biology and microbiota of trees are very different from those of herbaceous plants, so we cannot predict the behaviour of tree-microbiota interactions on the basis of what is known from herbaceous plants. While root exudates contribute to the initial steps of selection of the rhizospheric and root microbiota, plant metabolites also participate in the structuring of host endophytic microbiota (Reinhold-Hurek et al., 2015; Trivedi et al., 2020). For instance, it has been suggested that variations in the microbiome between poplar species are linked to specific differences in defence compounds, such as the biosynthesis of phenolic glycosides (salicylates) and other metabolites (Lindroth and St. Clair, 2013; Tschaplinski et al., 2014; Veach et al., 2020, 2019). Conversely, changes in the microbial composition of roots can significantly affect the metabolome of poplar roots and shoots (Mangeot-Peter et al., 2020).

In light of all of these knowledge gaps, we aimed in this study to characterize the *Populus tremula x tremuloides* T89 root exudates, the early dynamics of the assembly of the microbiota along the root-to-shoot axis, and its interaction with shoot and root metabolite contents. This study investigated the dynamics of microbial colonisation of roots and shoots of naive poplar cuttings from the soil reservoir combined with the dynamics of root exudate composition, as well as metabolomics of roots and shoots. Naive - i.e., entirely sterile at the time of planting - *Populus tremula x tremuloides* T89 cuttings were cultivated in either natural or sterilised (gamma-irradiated) soils for 30 days in small, closed mesocosms. Root and shoot biomass, root exudate composition, soil, rhizosphere, root and shoot microbiota over time, and root and shoot metabolomics at the final time-point were measured. We hypothesised that: (1) the presence of soil microbiota modifies metabolite contents (both composition and quantity) of root exudates; (2) the root exudation is a dynamic process in time and correlates with the assembly of specific microbial communities in the rhizosphere; (3) microbial colonisation of roots and shoots induces metabolomic changes of both roots and shoots; (4) the establishment of aboveground communities follows the same dynamics as belowground; and (5) specific microbial communities are selected from the soil reservoir to colonise the shoots.

## RESULTS

### 1. Root exudate composition is dynamic over time and microorganisms strongly reduce the abundance of root exudate metabolites

In order to investigate how microbial communities influence poplar root exudation, the metabolite profiles of root exudates were characterized over time in the presence or absence of microorganisms (**Figure S1**). First, the possible impacts of gamma-irradiation on soil fertility and growth of poplar cuttings were examined 30 days post-planting. Gamma irradiation of the soil did not have a significant impact on soil carbon or nitrogen levels, or on pH. Nor did it affect the availability of Ca, Fe, Mg, K, and Na, but gamma irradiation did reduce the levels of phosphorus in the soil by 0.3x times (**Table S1**). Shoots and roots grew similarly with or without microbes (**Figure S2**). Between 15 and 72 metabolites were detected in the root exudates of young poplar cuttings (**Figure 1A**). Eighty percent could be attributed to known metabolites belonging to five main classes of compounds: glycosides (23%), organic acids (13%), defence compounds (10%), lipid-related metabolites (7%), sugars (7%), and amino acids (2%) (**Figure 1B, Figure S3**). Striking differences in the metabolite composition of root exudates were observed between poplars grown in natural or sterilised soil. The number of metabolites detected in root exudates were 3x to 5x times higher in sterilised soil in comparison with natural soil (**Figure 1A**). Root exudates captured from poplars grown in sterilised soil were enriched in all major classes of metabolites, including defence compounds (e.g., phenylethyl-tremuloidin, salicyl alcohol, salicylic acid), organic acids (e.g., hexanoic acid, citric acid, ferulic acid), and sugars (e.g., glucose, sucrose, galactose) in comparison with root exudates from cuttings grown in natural soil (**Figure 1B**, **Figure 2, Figure S3, Figure S4**). Interestingly, most of the metabolites belonging to the glycoside, amino acid (e.g., 5-oxo-proline, GABA), and lipid-related (e.g., monopalmitin, monostearin, palmitic acid) classes were only detected in root exudates from poplars grown in sterilised soil (**Figure 1B, Figure S3).** The lipid related monopalmitin and monostearin were by far the most abundant compounds found in the root exudates of poplars grown under sterile conditions, being 10x and 15x times, respectively, more abundant than the most abundant sugars and organic acids **(Table S2)**. In addition, root exudation profiles were dynamic over the 30 days of poplar growth in both soil types but followed opposite trends. While a significant decrease of the number of metabolites of root exudates produced in natural soil was observed, this number increased significantly in sterilised soil over the 30 days (**Figure 1A**). In sterilised soil, the production of most root exudates, - belonging to diverse metabolite classes - increased significantly over time (e.g., Defence: tremuloidin, phenylethyl-tremuloidin; Organic acid: citric acid, erythronic acid; Lipid-related: monopalmitin, monostearin; Sugars: glucose, butyl-mannoside) (**Figure 1B**). Conversely, root exudates from poplars grown in natural soil displayed increased concentration of glycerol, whereas the concentration of an unidentified glycoside (14.21 min; m/z 279) and two unidentified compounds (12.06 min; m/z 404 517 307 319, 13.21 min; m/z 235 204 217) decreased significantly over time (**Figure S3**).

**Figure 1.**
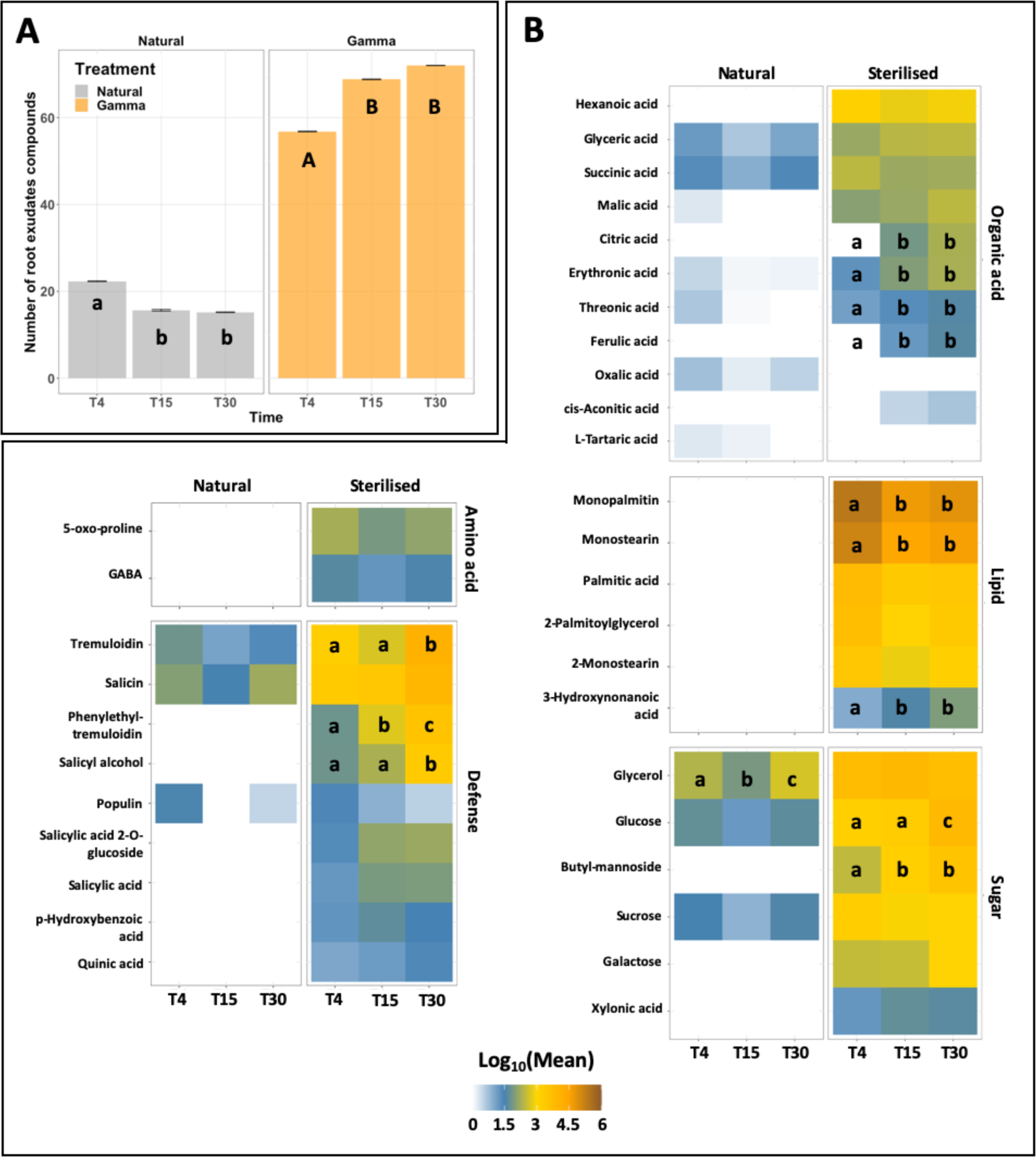
Influence of microorganisms on the root exudates profile over time. **(A)** Root exudates richness of poplar grown on natural and sterilised soil over 30 days of growth. **(B)** Dynamics of root exudates of poplar grown in presence or absence of microorganisms. Values correspond to the exudate mean concentration transformed by Log10. Letters indicate significant differences of metabolite concentration over time for each treatment (n = 5, Kruskal-Wallis, FDR corrections, p.adj ≤ 0.05, Fisher LSD post-hoc test).

**Figure 2.**
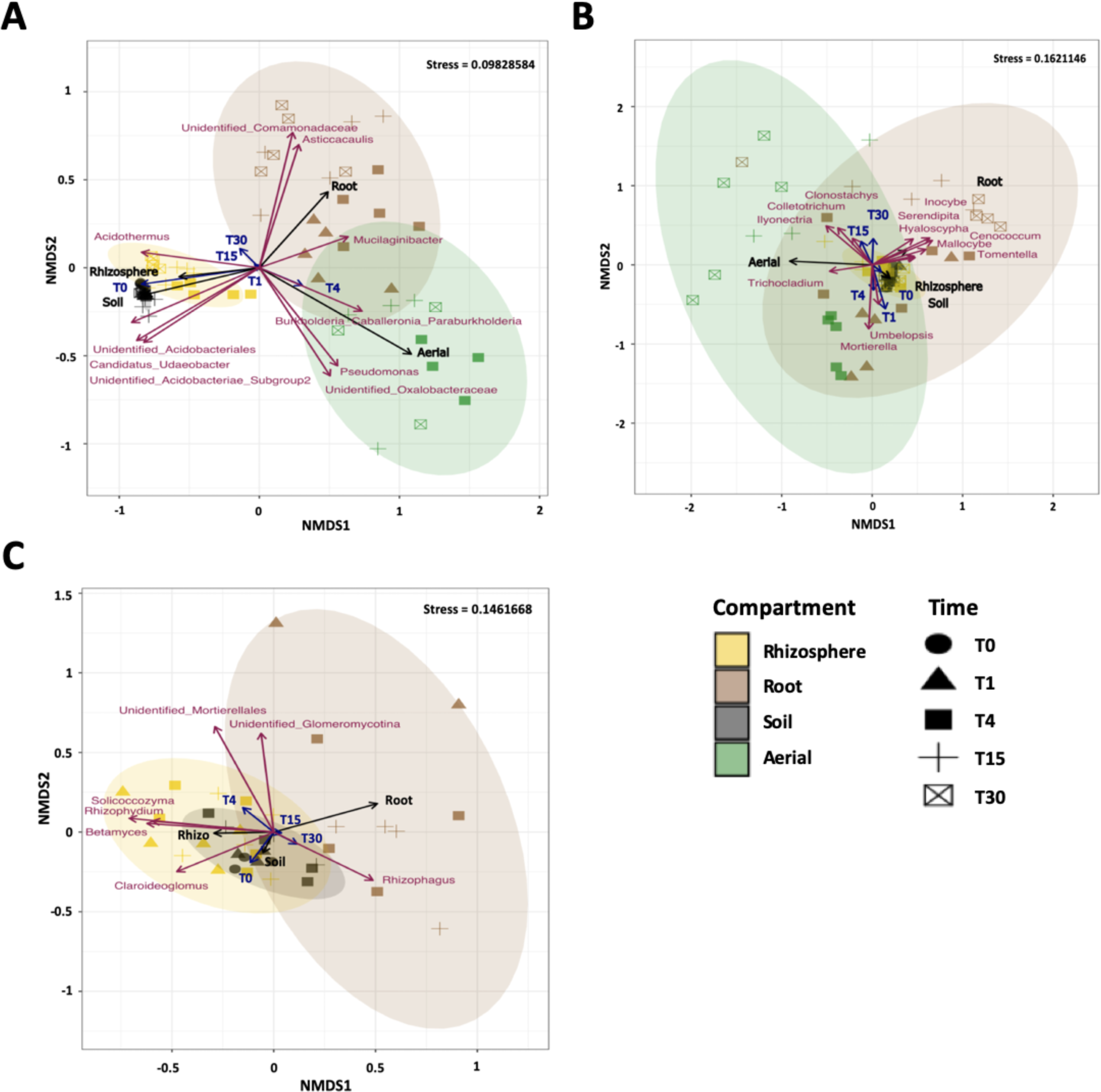
Structure of microbial communities in distinct compartments over 30 days of growth. NMDS representation of **(A)** bacterial, **(B)** fungal and **(C)** Glomerales communities over 30 days of growth. For all **A**, **B** and **C**, vectors indicate significant variables structuring the microbial communities after multiple regression analyses and 1,000 permutations (n = 3-5, FDR corrected, p.adj ≤ 0.01).

To conclude, root exudates of young poplar cuttings are dynamic over-time, from 4 to 30 days after planting, and root exudates of poplar cultivated in the presence of microorganisms contain less metabolites than in absence of microorganisms.

### 2. Microbial communities from the rhizosphere but not soil evolved over time

The massive alteration of root exudates in the presence of microorganisms suggests that the rhizospheric microbiota consume a large fraction of the exudates reducing their concentrations to below the level of detection and/or the existence of feedback effects of the microbiota on plant metabolism. In order to get a better understanding of the microorganisms involved in these processes, the fungal and bacterial communities of the rhizosphere and their dynamics were characterized.

Given that the soil was the main reservoir of microorganisms colonising poplar habitats in our experimental design, the microbial communities present in the soil before transplanting axenic poplars were characterized first. A total of 286 ± 6 fungal Operational Taxonomic Units (OTUs) and 941 ± 2 bacterial OTUs were detected in soil (**Table S3.A**). Fungal soil communities were dominated by endophytes, ectomycorrhizal fungi (EMF), and to a lesser extent saprotrophs (respectively 28 ± 1%, 26 ± 4%, 14 ± 1%) (**Figure S5**, **Table S3.B**). The endophyte *Mortierella*, and the EMF *Inocybe*, and *Tuber* were the most dominant fungal genera detected in soil over time (**Table S3.C**). Regarding arbuscular mycorrhiza fungal (AMF) communities that were tracked independently with 28S barcode sequencing, *Rhizophagus*, *Glomus* and an unidentified OTU of *Glomeromycetes* were the most abundant genera in soil (**Table S3.D**). Finally, Candidatus *Udaeobacter* (Verrucomicrobia) and two unidentified OTUs of the Acidobacteria phylum dominated soil bacterial communities over the 30 days of growth (**Table S3.E**). Overall, soil bacterial and fungal communities, -including Glomerales-, remained stable over time.

By contrast, the diversity and composition of fungal and bacterial communities fluctuated overtime in the rhizosphere, with the exception of AMF. After 4 days of growth, 255 ± 12 fungal OTUs were identified, which increased to 269 ± 7 by the end of the experiment. Similarly, the 950 ± 22 bacterial OTUs detected at the early time point, increased to 997 ± 5 after 30 days of growth (**Table S3.A**). The rhizospheric bacterial community was dominated by Proteobacteria (*Pseudomonas, Burkholderia, Oxalobacteraceae*), Verrucomicrobia (Candidatus *Udaeobacter*), Bacteroidetes (*Mucilaginibacter*) and Acidobacteria (Candidatus *Solibacter*) while EMF (e.g., *Inocybe, Lactarius, Tomentella, Tuber*), endophytes (e.g., *Mortierella, Hyaloscypha, Ilyonectria*) and to a lesser extent, saprophytes (*Umbelopsis*, *Bifiguratus*) dominated the fungal community (**Figure S6, Figure S7, Table S3.C-E**). Many of these microorganisms were enriched in the rhizosphere compared to soil, illustrating the well-known selective effect of this habitat, it is also noteworthy that Candidatus *Udaeobacter,* Candidatus *Solibacter* and *Acidothermus* were as abundant in the rhizosphere and soil, representing more than 15% of the reads in these two habitats. Furthermore, different dynamics of colonisation were observed in the rhizosphere among the dominant bacterial and fungal genera (>3%, p.adj ≤ 0.05). The relative abundance of most of the bacterial genera that were strongly enriched in the rhizosphere compared to soil, such as *Burkholderia, Pseudomonas* and *Mucilaginibacter*, decreased significantly over time, while members of Candidatus *Udaeobacter,* Candidatus *Solibacter* and *Acidothermus* remained stable from T4 to T30. Regarding fungi, despite no significant difference in the abundance of the main fungal trophic guilds over time (p.adj > 0.05), the relative abundances of the saprotroph *Bifiguratus* and the EMF *Inocybe* increased significantly, while the EMF *Lactarius* decreased over time. The fungal endophyte *Ilyonectria* was only significantly more abundant at T15, but not at T30 (**Figure S6, Figure S7, Table S3.C**). Lastly, as for the soil, *Glomus*, *Claroideoglomus* and *Rhizophagus* dominated the rhizosphere compartment and remained stable over the 30 days of growth.

Overall, while microbial communities in the soil remained stable over the experiment, the assembly of the communities belonging to the rhizosphere was dynamic over time, with some dominant fungal and bacterial genera found only transiently.

### 3. Microbial colonisation from belowground to above ground compartments

Among the dominant fungal and bacterial genera in the rhizosphere that have been only found transiently, microorganisms, such as *Pseudomonas* or *Ilyonectria*, are known to be potential root and leaf endophytes (Compant et al., 2021; Henning et al., 2016; Liao et al., 2019). We thus surmised whether this timely detection in the rhizosphere reflected a transitory movement towards their final habitat (root and/or shoot), or whether they were outcompeted by other microorganisms. To answer this question, the dynamics of the fungal and bacterial communities from the rhizosphere to the roots and the shoots were followed. Microbial colonisation was rapid and highly dynamic in both belowground and aboveground compartments (**Figure 2, Table S3**). Fungal and bacterial taxa in root systems were detected as soon as after 1 day of growth, and bacterial communities were already present in shoots, even though their relative abundance was variable among the samples, and thus, not considered in the analyses.

After only 4 days, both bacterial and fungal communities were established in roots and shoots. Fungal endophytes dominated both root and shoot fungal communities at the early time points and decreased over time. While EMF dominated the late stage of root colonisation, saprotrophs and pathogens were the most abundant fungal guilds detected in shoots after 30 days. In contrast, the dominant Glomerales, including *Glomus* and *Rhizophagus,* remained stable in roots over time (**Figure S6, Figure S7, Table S3.D).** As indicated by the analyses of microbial structure, early root and shoot fungal communities were closely related before differentiating over time (**Figure 2, Table S4**). The fungal endophyte *Mortierella* and the saprotroph *Umbelopsis* drove both root and shoot early fungal communities before vanishing from those compartments at later stages of colonisation (**Figure 2**, **Figure 3**).

**Figure 3.**
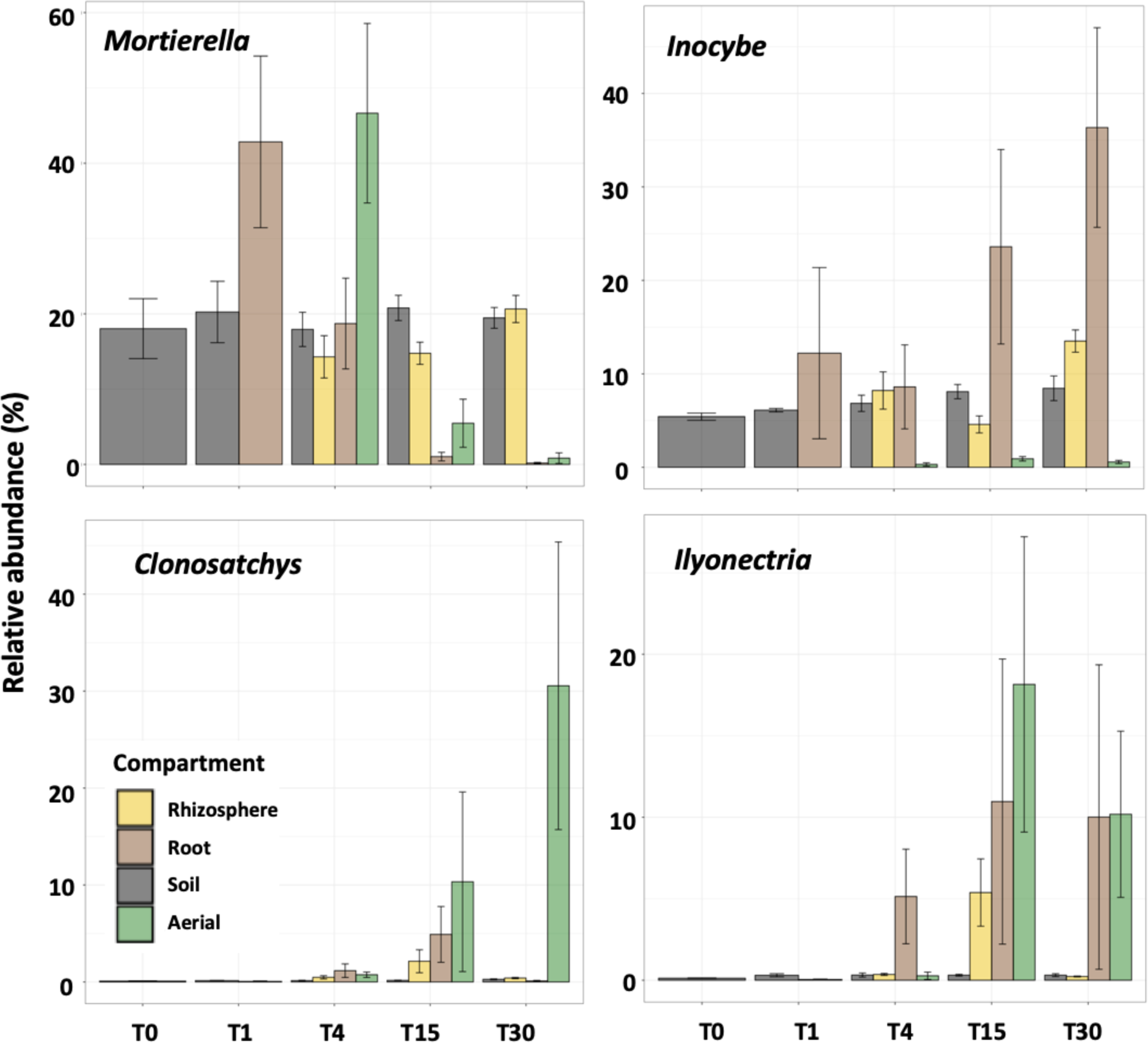
Relative abundance of fungal genera associated with specific habitat over 30 days of growth. Fungal taxa were chosen according to their significance related to particular time or habitat in multivariate partition analyses after multiple regression analyses and 1,000 permutation (FDR corrected, p.adj ≤ 0.01). Histograms represent the mean relative abundance of each taxa and bars indicate their standard error (n = 3-5).

These fungal genera were later replaced by specific taxa depending on the plant compartments. A core fungal microbiota was detected where some taxa assembled in both compartments, although other microorganisms were specific to a particular niche (**Figure S6, Figure S7**). For example, the endophyte *Ilyonectria* colonised both roots and shoots in similar relative abundances, whereas *Trichocladium*, *Colletotrichum* and *Clonostachys* dominated aboveground compartments, and the fungal endophyte *Hyaloscypha* and the EMF *Mallocybe*, *Inocybe*, and *Tomentella* prevailed in belowground compartments (**Figure 2**, **Figure 3, Figure S8**). It is noteworthy that these EMF were also detected in shoots, not being an isolated event, as they remained detectable at low levels until 30 days of growth. Even though this effect was less striking for bacteria than for fungal communities, the transfer of bacterial genera from belowground to aboveground compartments was also detected (**Figure 2**, **Figure 4, Figure S9**). *Mucilaginibacter*, *Pseudomonas*, and *Burkholderia-Caballeronia-Paraburkholderia* were present in root systems at an early stage of colonisation before prevailing in aerial compartments, while *Asticcacaulis*, an unidentified OTU of the Comamonadaceae family, and *Acidothermus* dominated belowground compartments (**Figure 2**, **Figure 4, Figure S9**).

**Figure 4.**
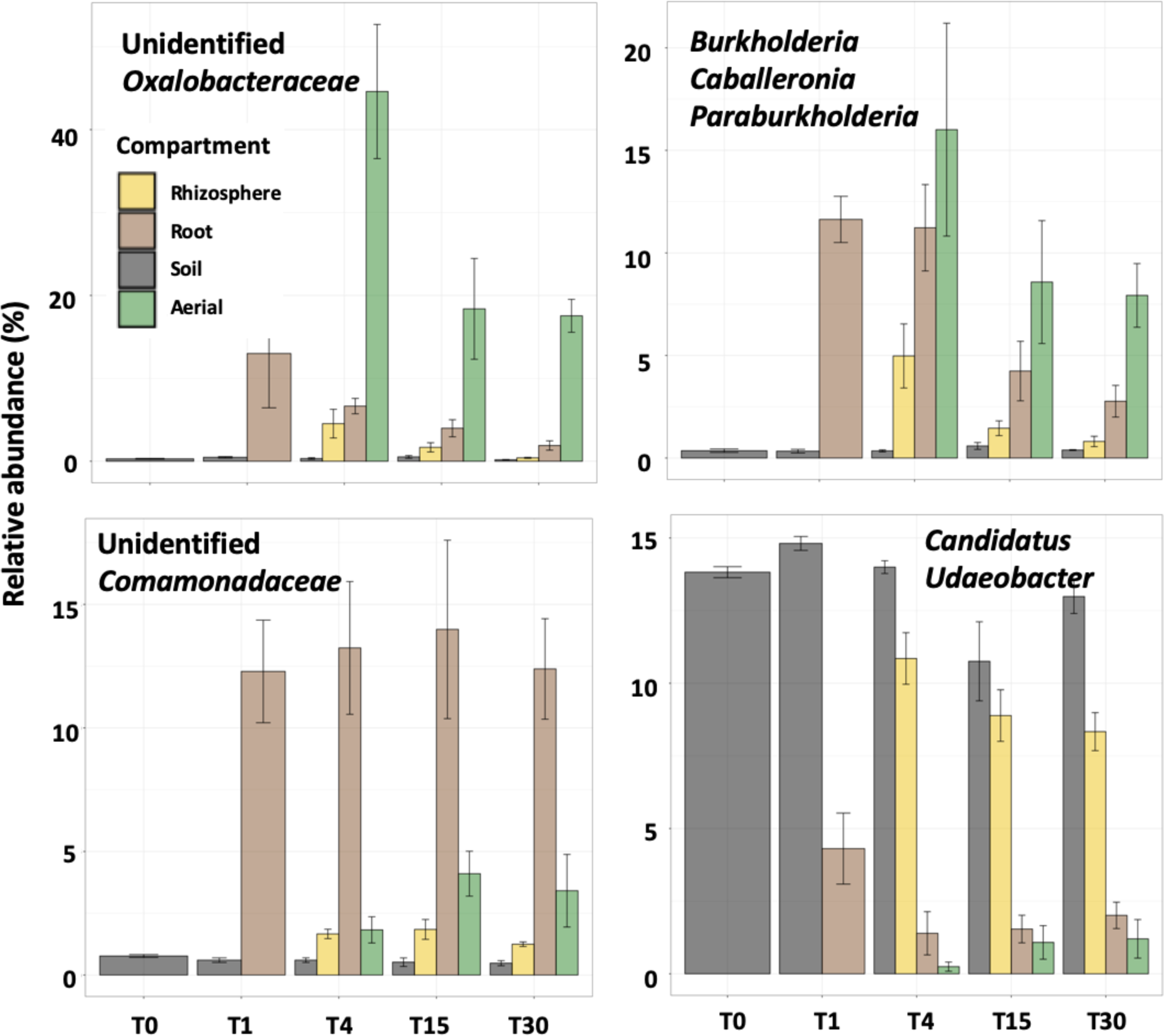
Relative abundance of bacterial genera associated with specific habitat over 30 days of growth. Bacterial taxa were chosen according to their significance related to particular time or habitat in multivariate partition analyses after multiple regression analyses and 1,000 permutation (FDR corrected, p.adj ≤ 0.01). Histograms represent the mean relative abundance of each taxa and bars indicate their standard error (n = 3-5).

To conclude, the microbial colonisation of the root and shoot habitats evolved over time, through the transition of a core and a specific microbiota from below to aboveground compartments. EMF dominated root systems while saprotrophs and pathogens dominated the shoots.

### 4. Microbial communities alter belowground and aboveground poplar metabolite composition

Having demonstrated that root exudates were strongly impacted by microbial presence and that both roots and shoots were massively colonised by complex, dynamic and specific microbial communities, we investigated how microbial colonisation influenced the root and shoot metabolomes. The metabolomic profiles of belowground and aboveground compartments were characterized after 30 days of growth in the presence or absence of microorganisms and correlated metabolic profiles with microbial communities.

Similar to root exudate responses, metabolite richness and diversity were higher in poplars grown in sterilised soil after 30 days in comparison with natural soil, particularly in roots. For poplars grown in natural soil, 64 and 90 metabolites in belowground and aboveground compartments were detected respectively, which increased to 81 and 96 metabolites, respectively, in sterilised soil (**Figure 5, Figure S4**).

**Figure 5.**
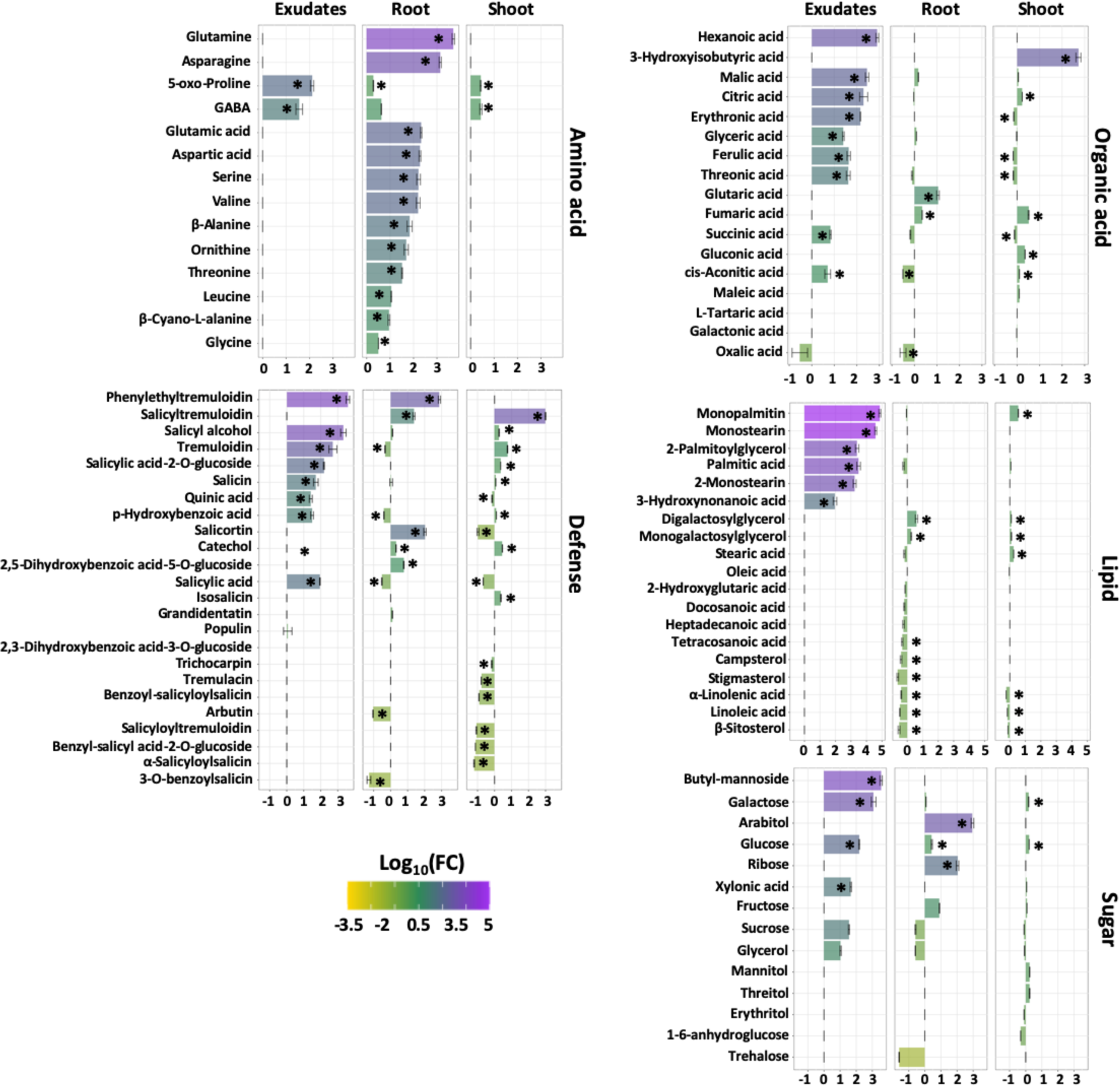
Influence of microorganisms on the composition and abundance of root exudates, root and shoot metabolites after 30 days of growth. Bars represent the log2 fold change of the relative abundance of metabolic compounds detected in sterilised soil (positive bars) versus natural soil (negative bars). * indicate significant difference of metabolite abundance between the two treatments (n = 5-25, Wilcoxon, FDR corrections, p.adj ≤ 0.05).

Microbial colonisation induced greater variation in metabolite concentrations in the roots than the shoots (**Figure 5, Figure S4**). Most striking was the reduction of the levels of most amino acids, as well as glucose and fructose, in roots in the presence of microorganisms. By contrast, levels of glycerol, sucrose, and trehalose increased in roots of poplars grown in natural soil. Microbial colonisation also led to an increase of sterol levels in roots and of the unsaturated fatty acids, α-linolenic, and linoleic acid in both roots and shoots (**Figure 5**). Interestingly, the majority of defence metabolites and their conjugates were detected in the aerial compartments (e.g., trichocarpin, tremulacin, salicyltremuloidin) and varied differently depending on plant organ (**Figure 5**). Tremuloidin decreased significantly in root systems from sterile soil, but it increased in shoots of poplars grown in the same soil (**Figure 5**). Surprisingly, salicylic acid and salicyltremuloidin were more readily detected in both roots and shoots of poplars grown in absence of microbes (**Figure 5**).

Overall, our data show that as early as 30 days post-planting, root and shoot metabolomes of naive poplar cuttings are strongly modified by root microbial communities.

### 5. Correlations between poplar metabolites and microbial taxa abundances

After showing that microorganisms alter the metabolite profiles of poplar in both belowground and aboveground habitats, we investigated whether the presence of particular microbial communities was associated with specific metabolites in root exudates, roots and shoots using multiple regressions models through Redundancy Analyses (RDA).

Significant correlations between root exudates and microbial communities were observed over time (**Figure 6**). Although only a novel metabolite, tentatively identified as 2-*o*-Benzoyl-*p*-toluic acid glucoside, was positively associated with the fungi *Mortierella* and *Inocybe*, 4 compounds (glycerol, L-tartaric acid, glyceric acid, and the unidentified 13.56 min; m/z 273 363) were correlated with bacterial genera (**Figure 6.A**). The relative abundance of early associated *Pseudomonas* (Gamma Proteobacteria), *Pedobacter* (Bacteroidetes), *Burkholderia* and *Cupriavidus* (Beta Proteobacteria), *Rhizobium* (Alpha Proteobacteria), and *Mucilaginibacter* (Bacteroidetes) were positively correlated with the levels of two organic acids, L-tartaric acid and glyceric acid. In contrast, the levels of late bacterial taxa were more (e.g., *Ktedonobacteraceae JG30a* and *Bryobacter*) or less associated with glycerol (eg., *Bdellovibrio*), respectively which was enriched at the end of the experiment (**Figure 6.B**).

**Figure 6.**
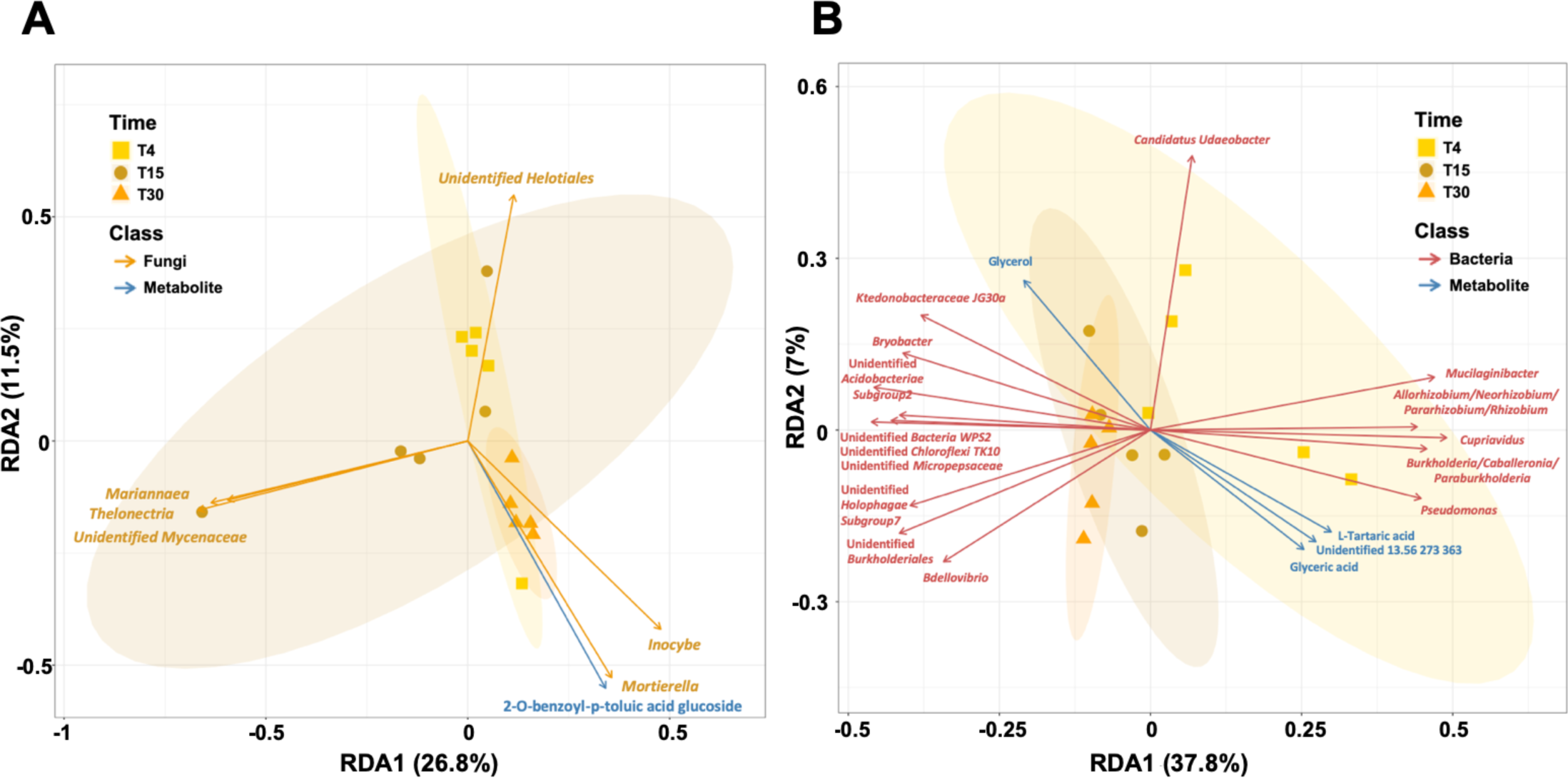
Redundancy analysis showing the correlations between (A) fungal or (B) bacterial taxa and root exudates in the rhizosphere over 30 days of growth. Vectors indicate significant variables structuring the microbial communities after multiple regression analyses and 1,000 permutations (n = 3-5, FDR corrected, p.adj ≤ 0.01).

Within plant tissues, RDA analyses revealed associations between 6 fungal taxa and 16 metabolites (**Figure 7.A**), and 14 bacterial taxa and 27 metabolites (**Figure 7.B**). Associations between shoot metabolites and microbes tended to be more numerous than those between root metabolites and microbes. Four metabolites - the sugar acid/alcohol, xylonic acid and threitol, the defence compound salicylic acid, and the antioxidant alpha-tocopherol - were positively correlated with fungal and bacterial taxa in shoots (**Figure 7.A-B**). The EMF *Inocybe*, the endophyte *Hyaloscypha*, and the saprophyte *Luellia* that all mainly colonized roots, were positively associated with 4 metabolites that were only detected in the roots at T30, including the lipid-related metabolite tetracosanoic acid, the two glycosides purpurein and grandidentatin, and 2-hydroxyglutaconic acid (**Figure 7.A**). Additionally, these fungi were significantly positively correlated with sucrose. Conversely, the shoot-associated fungi, *Clonostachys*, *Bifiguratusi*, and *Trichocladium*, were positively associated with several metabolites that were enriched in the shoots, including the defence metabolite salicylic acid, sugar alcohol/acid (threitol, xylonic acid), organic acids (erythronic acid, maleic acid), lignin precursor (caffeic acid), and several other compounds of unknown function (e.g., *o*-cresol glucoside…) (**Figure 7.A**). Regarding bacteria, only one uncharacterized metabolite (9.60 min; m/z 228 110 291) was found associated with bacterial taxa in roots (**Figure 7.B**). In contrast, 10 bacterial genera that were enriched in shoot tissues were associated at different degrees with shoot metabolites. The strongest associations were found for OTUs of the Oxalobacteraceae and Micrococcaceae families, and the genera *Mucilaginibacter* (Bacteroidetes) and *Catenulispora* (Actinomycetes) with several defence compounds (salirepin, salicylic acid, tremulacin), organic acids (malic acid, aconitric acid, galactonic acid) and several glucosides (**Figure 7.B**). Those compounds were also less strongly associated with bacteria belonging to *Dyella*, *Pseudomonas* (Gamma Proteobacteria), *Pedobacter* (Bacteroidetes), *Burkholderia* (Beta Proteobacteria) and *Rhizobium* (Alpha Proteobacteria) (**Figure 7.B**).

**Figure 7.**
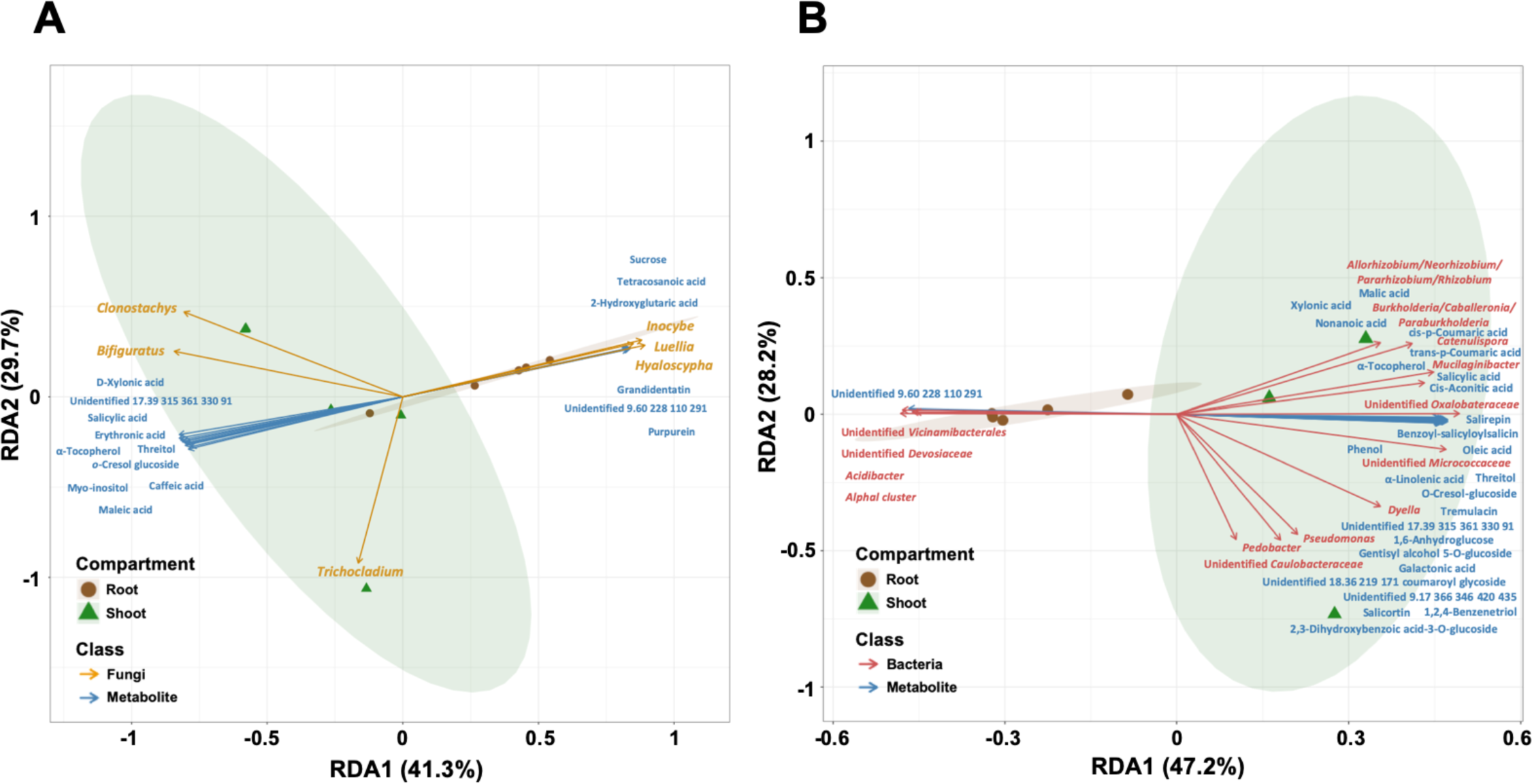
Redundancy analysis showing the correlations between (A) fungal or (B) bacterial taxa and metabolite compounds in poplar root and shoot systems after 30 days of growth. Vectors indicate significant variables structuring the microbial communities after multiple regression analyses and 1,000 permutations (n = 3-5, FDR corrected, p.adj ≤ 0.01, bootstrap > 90%).

Overall, these RDA analyses revealed metabolite and microbial biomarkers in root and shoot tissues, with specific metabolites highly enriched in either tissue.

## DISCUSSION

It is now well demonstrated that nearly all tissues of plants harbor microbial communities and that plants, including poplars, offer different habitats selecting a contrasting microbiota, particularly between roots and shoots (Beckers et al., 2017; Bettenfeld et al., 2021; Brown et al., 2020; Cregger et al., 2018; Maignien et al., 2014; Wallace et al., 2018; Wei et al., 2021). Extensive research has been carried out to determine the factors controlling the structuring of rhizosphere and root microbial communities in a wide range of plants, from herbaceous plants to trees (Trivedi et al., 2020). In *Populus* sp., like many plant species, soil is the major driver determining the rhizosphere microbiota (Beckers et al., 2017; Bonito et al., 2014; Mangeot-Peter et al., 2020). Rhizodeposition and factors dependent on the host tree, such as immunity, and for poplar, salicylate-related compounds, are thought to refine the selection of the microbiota in the rhizosphere (Veach et al., 2019). However, most studies are based on comparing microbiota between tissues at a given time, and less is known about the process of colonisation and selection of microbiota between tissues, particularly for perennials (Dove et al., 2021; Edwards et al., 2018; Fracchia et al., 2021). Here, the very early dynamic nature of this process was investigated to capture the different stages of the colonisation by fungi and bacteria of belowground and aboveground tissues. We combined these data with the analyses of the composition of root exudates of 1-month-old *Populus tremula x tremuloides* T89 cuttings over 30 days and of root and shoot metabolomes at 30 days. Evidence indicate that (1) the presence of microbiota massively modifies the composition of root exudate, and root and shoot metabolomes, (2) root exudation is dynamic over time, as is the microbiota of the rhizosphere, roots, and shoots, (3) soil is a reservoir of microorganisms for the colonisation of shoots, and (4) roots and shoots are first colonised by the same microorganisms that are later replaced by habitat-specific taxa.

### Composition of *Populus tremula x tremuloides* root exudates from young cuttings

Limited information exists regarding the composition of poplar root exudates. A study by Li et al., (2022) endeavored to analyse the root exudates of four poplar species, establishing correlations between poplar root exudate metabolites and the predominant bacterial taxa in the rhizosphere. However, this study primarily focused on collecting rhizospheric soil metabolomes rather than authentic root exudates, complicating direct comparisons with our research. In our study, we investigated the root exudates of *Populus tremula x tremuloides* T89, revealing a rich composition encompassing sugars, organic acids, glucosides, and various phenolic compounds, including flavonoids and lipid-related metabolites. This composition aligns with existing research on non-perennial species, confirming poplar root exudates as a carbon-rich environment, which likely serve as nutrients to feed soil microbes (Sasse et al., 2018) and then participate in the soil C-cycle (Dijkstra et al., 2021). As well as being a source of nutrients, root exudates also contain secondary metabolites, notably phenolic compounds, capable of regulating the growth of microorganisms (Pang et al., 2021). More specifically to poplars, salicylates (tremuloidin, salicin, populin, salicylic acid) and their derivatives were identified within the root exudates. Although salicylic acid (SA) has been detected in the root exudates of several non-perennial species (Khorassani et al., 2011; Zhalnina et al., 2018), concentrations and diversity of salicylates tend to be lower in the root exudates of other plants. These compounds exhibit multifaceted functions in root exudates acting as deterrents at low concentrations for microorganisms (Lebeis et al., 2015), while attracting saprotrophs capable of breaking them down (Clocchiatti et al., 2021), or participating in phosphate solubilization in soil (Khorassani et al., 2011). In *Arabidopsis*, the abundance of some bacterial communities is impacted in response to SA signalling, and it is partly explained, in part, by the use of SA as a C source for bacterial growth or as an immune signal (Lebeis et al., 2015). Clocchiatti et al., (2021) demonstrated that the combination of SA acid and primary metabolites induces a shift in the balance between fungi and bacteria, favoring the growth of saprotrophic fungal. Finding the same compounds in poplar root exudates led us to propose that salicylate compounds not only play a role for selection of endosphere microbiota, as notably suggested by Veach et al., (2019), but also contribute to the selection of microbes from the soil reservoir.

### Lipid-related metabolites involved in root-microbe signaling versus sustaining endophytic microbial growth

Interestingly, our study indicated a lipid-related signature for poplar root exudates and roots. Whereas mainly fatty acids were detected in poplar root exudates, sterols and specific fatty acids were also identified in poplar roots. Fatty acids in root exudates were specifically detected from poplars cultivated in absence of microbes. This is consistent with a possible role of these fatty acids in plant root-microbe signaling (Macabuhay et al., 2022). Fatty acids from *Pinus sylvestris* root exudates have been previously shown to stimulate the growth of the EMF *Laccaria* and *Leccinum* (Fries et al., 1985). Additionally, the observed consumption of fatty acids at 30 days is consistent with the detected presence of arbuscular mycorrhizal fungi (*Rhizophagus*, *Glomus* OTUs). Indeed, several studies demonstrated lipid transfer from the plant to the AM-fungi in the form of monoacylglycerol containing C16-fatty acids (Bravo et al., 2017; Jiang et al., 2017; Keymer et al., 2017; Luginbuehl et al., 2017). In contrast, root microbial colonisation induced higher concentration of phytosterols (campsterol; stigmasterol; β-sitosterol) and fatty acids (stearic acid, α-linoleic acid, linoleic acid) in roots. These accumulations likely support the demand for membrane remodelling to sustain microbial colonisation, in particular for arbuscule formation by AMF, but also for other types of endophytic fungi (Macabuhay et al., 2022; Roth et al., 2019).

### Reduction in root exudates: active consumption by microbes or negative feedback?

Massive differences between root exudates from poplar trees grown in the presence or absence of microbes were detected as early as 4 days after planting. Lipid-related metabolites, sugars, organic acids, amino acids, salicylates and derivatives were greatly depleted in presence of microbes as early as 4 days post plantation. At this time-point, mainly bacteria and saprotrophic fungi were colonising the rhizosphere and the roots, which argues in favour of an important role of bacteria and saprotrophic fungi in root exudates consumption, including fatty acids and defence compounds. The degradation of these defence compounds may later facilitate the development of sensitive microorganisms that could not have developed in the presence of phenolics (Clocchiatti et al., 2021). At later time points (T15 and T30), the same trend for metabolite depletion of root exudates in presence of microbes was found. The question remains whether soil microbes consumed poplar root exudates as nutrients sources or to degrade toxic defence compounds, allowing the entry of endophytic microbes. Whereas local microbial communities can trigger systemic changes in root exudation (Korenblum et al., 2020), we were not able to detect such alteration with our protocol of collecting root exudates from the whole root system. Yet, the presence of organic acids, such as glyceric acid and tartaric acid, were positively correlated with the relative abundance of early-stage bacterial taxa including *Pseudomonas*, *Burkholderia*, and *Mucilaginibacter*. As organic acids can act as chemoattractants for bacteria, these molecules could also serve as primary selection factors for microbes from the rhizosphere (De Weert et al., 2002; Feng et al., 2019; Zhang et al., 2014). Interestingly, late-stage bacterial taxa (e.g. *Ktedonobacteraceae JG30a*, *Bryobacter*) were positively correlated with glycerol but negatively correlated with glyceric acid. Given that glyceric acid is primarily derived from the microbial oxidation of glycerol (Habe et al., 2009), metabolite turnover represents a selection process either acting as a repellent or attractant depending on the bacterial taxa.

As expected, poplar root exudates contained low-molecular-weight carboxylates, such as malic acid, citric acid, and aconitic acid. These compounds were enriched in root exudates of poplars cultivated in sterilised soils. This is consistent with the mechanism of plants increasing P-uptake by secreting carboxylates that can displace immobilized P from inorganic and organic soil compounds (Bolan et al., 1994; Chai and Schachtman, 2022). The levels of these carboxylates were lower in root exudates from poplars cultivated in the presence of microbes, whereas the amount of oxalic acid was higher in roots of the same poplars. Oxalic acid production may also be of fungal origin. It could serve as a signal molecule for the mycophagous bacterium *Collimonas*, which has been detected in the roots of poplar in isolated cases (Rudnick et al., 2015). Taken together, these data suggest that P-mobilization by plant-produced organic acids is mainly occurring in absence of microbes associated with the root systems. It can be hypothesised that organic acids have a dual role: P-scavenging in absence of microbes and a C source for microbes.

### Modification of root and shoot metabolites as indicators for microbial community establishment

In contrast with poplar root exudates, metabolites accumulated in roots and shoots of poplars cultivated in presence of microbes rather than in their absence at 30 days post-plantation. Whereas root microbial colonisation induced higher sucrose and P concentrations (but less glucose and fructose) in roots, amino acids accumulated to high concentrations in roots of plants cultivated in sterilised soils. At that time-point, roots systems were mainly colonised by EMF, known to transfer N to the plant and to receive C in return (Bogar et al., 2022; Martin et al., 2016). These data support C exchange from sucrose breakdown to the microbes. Interestingly, sucrose was positively correlated with the presence of three fungi in the root: the EMF *Inocybe*, the endophyte *Hyaloscypha* and a microbe described as a saprotroph, suggesting that they may be greater consumers of sucrose produced by the plant. On the other hand, the lower concentrations of amino-acids in roots colonised by microbes were not expected. It can be hypothesised that amino acids are directly assimilated into proteins to sustain the increased metabolic processes in the presence of microbes, explaining the lack of amino-acids in our samples. Root microbial colonisation also strongly remodeled the poplar leaf defence compounds, as previously shown in other studies (Kaling et al., 2018). The levels of defence compounds were mainly correlated with the bacterial genera unlike fungi, suggesting that the modified niche (by metabolite changes) could be a trigger for the selection of bacteria colonising the leaves from the root system. Alternatively, bacteria may be the main trigger for the remodeling of leaf defence. Colonisation of the poplar tissues by soil-borne microorganisms was very rapid, with both fungi and bacteria being detected on the roots as early as 24 hours after the poplars were planted, and in the shoots shortly thereafter. The structuring of the microbial communities followed a two-step process for both root and aerial tissues in which an early-stage community, dominated by endophytes and saprophytes, rapidly colonised the tissues and was later replaced by a more stable community of symbionts. While the early-stage community of early colonisers was quite similar between roots and shoots, the late-stage communities were clearly differentiated between the roots and the shoots. However, dominant members of the shoot microbiota at the late time point were transiently detected at earlier time points in the rhizosphere and in the roots, suggesting that their first chemoattraction was in the rhizosphere and then followed by their transit through the roots to the shoots. It is noteworthy that levels of tartaric acid and glyceric acid in the root exudates followed the same trend of a decline over time similar to the bacterial taxa that were transiently detected, suggesting that they may act as chemoattractants in the rhizosphere.

Two main horizontal routes of colonisation can be envisaged for the phyllosphere: airborne microorganisms and those from insect carriers that land on leaves and form the epiphytic microbiota, including microorganisms that penetrate the leaf endosphere through stomata and wounds, versus microorganisms that travel from the soil through the roots and to the stems (Chaudhry et al., 2021), but the relative importance of the two routes is uncertain. Our data suggest that the soil may be an important reservoir of microorganisms for the colonisation of aerial tissues of *P. tremula x tremuloides* by both fungi and bacteria. This is in agreement with previous studies on grapes (Bettenfeld et al., 2021; Zarraonaindia et al., 2015), *A. thaliana* (Bai et al., 2015), and rice (Chi et al., 2005). The mesocosm device used in this study restricted the source of microorganisms that can colonise the phyllosphere to the soil, and it remains to be determined how significant is the airborne route in counteracting the soil reservoir. Nevertheless, the dominant taxa found in the shoots in our study including the fungus, *Ilyonectria* and the bacterial genera, *Pseudomonas, Burkholderia* and *Mucilaginibacter*. These microbes are typically found in the phyllosphere of various trees and plants (Cregger et al., 2018; Liao et al., 2019; Orozco-Mosqueda and Santoyo, 2021; Toju et al., 2019), suggesting that our observations are not an artefact and that these microorganisms can colonise shoot tissues from the soil. However, it remains to be determined whether they migrate to the shoots via the surface (epiphytic) or within the tissues (endophytic). The colonisation of roots and shoots in two waves is reminiscent of what we have recently described for roots of *P. tremula x alba* 717-1B4 (Fracchia et al., 2021). In both studies, an early, massive colonisation of the roots by the endophyte *Mortierella* was observed but in contrast with our previous experiment, the saprophytes *Umbelopsis* and *Saitozoma*, while being abundant in the soil and in the rhizosphere, did not colonise the roots of *P. tremula x tremuloides* T89, suggesting potential genotype-specific responses. Nevertheless, the replacement of *Mortierella* by other endophytes and EMF in both poplar species, and in both roots and shoots over time, is noteworthy. It may be hypothesised that fast-growing species such as *Mortierella* are quicker to colonise the host, but then compete with niche specialists such as EMF and endophytes, or are excluded by the host. However, *Mortierella* has been regularly isolated as a poplar endophyte and has even been shown to have plant growth-promoting properties (Liao et al., 2019), suggesting that it has the ability to establish in poplar tissues. Alternatively, the fungus may remain in the tissues but at a low level of abundance compared to other fungi and thus stay hidden until the death of the tissues where it is also often detected in the early stage of decay (Voříšková and Baldrian, 2013). Specific monitoring using quantitative PCR and metatranscriptomics would be necessary to elucidate the behavior of this ubiquitous fungus.

### The peculiar case of AMF as a stable community over time

Unlike other types of fungi, the composition of the AMF community remained stable over the course of the experiment once established in the roots. We previously demonstrated using Confocal Laser Scanning Microscopy that AMF establish symbiotic associations with *P. tremula x alba* 717-1B4 roots within 10 days, but we were unable to definitely identify the fungal species by metabarcoding (Fracchia et al., 2021). Bonito et al., (2014) also reported that classical ITS and 18S metabarcoding methods were not able to characterize the Glomeromycete community in poplar roots although these fungi are well known to colonise poplar roots (Baum and Makeschin; Chifflot et al., 2009; Gehring et al., 2006; Neville et al., 2002). To circumvent this problem, the nested PCR method developed by Brígido et al., (2017) was used and captured in detail the composition of AMF communities in roots of *P. tremula x tremuloides* T89, for the first time using high throughput sequencing. We demonstrate that several species belonging to the genera *Rhizophagus* and *Glomus* can colonise a single root system at the same time, unlike *Acaulospora* and *Claroideoglomus* that were only retrieved from soil. Such a pattern is in accordance with previous studies using regular Sanger sequencing identification methods (Beauchamp et al., 2006; Chifflot et al., 2009). It is generally considered that AMF dominate in roots at the juvenile stage of life of poplars and they are then replaced by EMF (Lopez-Aguillon R., 1989), and that environmental factors can influence the balance between EMF and AM (Gehring et al., 2006; Han et al., 2023). Our data indicate that AM and EMF can together colonise naive root systems and coexist, even when the EMF strongly expand.

## CONCLUSION

In this work, we showed that microbial colonisation triggered rapid and massive changes in the quality and quantity of poplar root exudates and led to a strong alteration of the root and shoot metabolomes. Furthermore, we demonstrated that the assembly of microbial communities in both belowground and aboveground habitats is highly dynamic involving successional waves of colonization. Our investigation reveals a close relationship between fungal communities establishing in the roots and shoots during the early stages of colonization, with subsequent differentiation in the later stages. These findings support the transition of microorganisms from below to the aboveground compartments, followed by the fine-tuned selection of the host resulting in the assembly of specific communities among the plant habitats, although we observed the presence of a core microbiota colonizing both niches. Poplars are unique among temperate forest trees, firstly because of their particular metabolism of salicylates, and secondly because of the double colonisation of their roots by AMF and EMF and the high abundance of endophytes in their roots. It would therefore be very interesting in the future to determine whether our results apply only to the Salicaceae family or whether they are more generic to trees.

## MATERIALS AND METHODS

### Biological material

In order to decipher the dynamics of poplar microbiota establishment of fungal and bacterial communities between aboveground and belowground compartments, poplar, *Populus tremula* x *tremuloides* T89, was cultivated *in vitro* sterile conditions on Murashige and Shoog (MS) (2.2g MS salts including vitamins, Duchefa; 0.8% Phytagel and 2% sucrose). Poplar cuttings were cultivated at 24°C in a growth chamber (photoperiod, 16h day; light intensity, 150 μmol.m^-2^.s^-1^) on MS supplemented with indole-3-butyric acid (IBA) (2 mg.L^-1^) for 1 week before being transferred on MS for 2 weeks until root development. This development growth protocol was used for all experiments.

### Soil collection & sterilisation by Gamma-irradiation

To obtain a forest-like microbial inoculum, the top soil horizon (0 to 20 cm) of a *Populus trichocarpa* x *deltoides* plantation was collected over an area of 1 m^2^ under 5 different trees. Soil was dried at room temperature and sieved at 2 mm diameter pore size before being further used. Three subsets of 20 g, that was stored at -80°C until further soil physico-chemical property analyses. In order to decipher the influence of microbial communities on poplar root exudates and metabolomes, a subset of 50 kg of soil was sterilizsd by gamma irradiation (45-65 kGy, Ionisos, France). The soil was packaged in individual plastic bags containing 200 g of soil prior to gamma irradiation. The sterilised (gamma-irradiated) soil was stored for 3 months at room temperature in the dark before being used to allow outgassing of potentially toxic volatile compounds.

### Soil physico-chemical properties

Soil analyses were performed using the LAS (Laboratoire d’Analyses des Sols) technical platform of soil analyses at INRAe Arras, according to standard procedures, detailed online (https://www6.hautsdefrance.inra.fr/las/Methodes-d-analyse). Exchangeable cations were extracted in either 1M KCl (magnesium, calcium, sodium, iron, manganese) or 1M NH_4_Cl (potassium) and determined by ICP-AES (JY180 ULTRACE). The 1M KCl extract was also titrated using an automatic titrimeter (Mettler TS2DL25) to assess exchangeable H^+^ and aluminum cations (Al^3+^). Total carbon, nitrogen and phosphorus contents were obtained after combustion at 1000 °C and were determined using a Thermo Quest Type NCS 2500 analyser. The pH of the soil samples was measured in water at a soil to solution ratio of 1:2 (pH meter Mettler TSDL25). Exchangeable acidity was calculated by taking the sum of H^+^ and Al^3+^. The cation-exchange capacity (CEC) was calculated by taking the sum of both extracted exchangeable base cations and exchangeable acidity. Results are compiled in Table S1.

### Plant growth and sampling procedure

To investigate the dynamics of colonization of naive poplars by microbial communities, 200 g of soil, either natural or sterilised, were distributed into 1,500 cm^3^ boxes closed with filtered lids (OS 140 Box, Duchefa-biochimie), and soil was maintained at 75% humidity. Two uniform *in vitro* seedlings (1 cm long for shoots and 1-2 cm long roots) were transferred to pots containing the environmental soil, described above. Each pot was enclosed with a filtered cover allowing gas exchange, and the bottom was covered (approximately 1/3 of the pot) with aluminum foil to prevent algal and moss development. Plants were cultivated at 24 °C in a growth chamber under the same conditions described above (photoperiod, 16h day; light intensity, 150 μmol.m^-2^.s^-1^). In total, 100 plants distributed among 50 pots were grown over 1, 4, 15 and 30 days.

Regarding microbial community analyses, at each time point, bulk soil, rhizosphere (except at T1, where no adherent soil was observed), root and shoot samples from 5 plants were collected. The shoots and roots were separated and weighed, and the rhizosphere was collected by pouring the root systems with adherent soil in 15 mL falcon tubes containing 2 mL sterile 1X phosphate-buffered saline (PBS: 0.13 M NaCl, 7mM Na_2_HPO_4_, 3mM NaH_2_PO_4_ [pH 7.2]). After removing the root systems, the samples were briefly vortexed in the falcon tubes containing the rhizosphere and centrifuged for 10 minutes at 4000 rpm. Then, the supernatant was removed to only retain the rhizosphere samples. Finally, the roots were washed in sterile water to remove remaining soil particles. Soil, rhizosphere, shoot and root samples were frozen in liquid nitrogen and stored at -80 °C until DNA extraction. *In vitro* poplars were also harvested to confirm their axenic status prior to planting (time point T0).

We analysed the metabolite composition for both shoot and root habitats after 30 days of growth by harvesting 25 seedlings. Shoots and roots corresponding to each poplar line were pooled to obtain between 9 to 15 replicates of dry material ranging between 25 to 100 mg for both organs. The dry shoot and root material were ground using metal beads and tissue-lyzer before sample extraction and analysis of their metabolomic composition by GC-MS.

In addition, we followed the exudates composition from 4 to 30 days of growth. Root systems of plantlets were left to exude in sterile water for 4 hours and root exudates were collected for GC-MS analysis after 0.2µm filtering. Root exudates were purified using Sep-Pak C18 cartridges (Waters^TM^) in order to remove salts contained in the hydroponic solution. Briefly, the column was conditioned by loading 700μl (one volume) 7 times with 100% acetonitrile. The column was then equilibrated with 7 volumes of H_2_O before loading 2ml of exudate. The columns were washed with 5 volumes of water and eluted in three steps; with an acetonitrile gradient ranging from 20%, 50%, and 100%. Finally, the lyophilised root exudates were weighed, and their metabolomic profile analysed by GC-MS.

### Microbial community analyses

To investigate the establishment of microbial communities in distinct organs of axenic poplar *Populus tremula* x *tremuloides* T89, bulk soil, rhizosphere, roots and shoots were sampled after 1, 4, 15 and 30 days of growth. For soil and rhizosphere, DNA was extracted from 250 mg of material using DNeasy PowerSoil kit using the protocol provided by the manufacturer (Qiagen). For root and shoot samples, 50 mg of ground plant material (less than 50mg for root systems at T0, T1, and T4) were used to extract DNA using DNeasy Plant Mini kit following the manufacturer protocol (Qiagen). DNA concentration was quantified using a NanoDrop 1000 spectrophotometer (NanoDrop Products) and DNA extraction was normalised to the final concentration of 10 ng.µL^-1^ for soil and rhizosphere samples and 5 ng.µL^-1^ for root and shoot samples. To maximise the coverage of bacterial 16S rRNA and fungal ITS2 rRNA regions, a mix of forward and reverse primers was used as previously described (Fracchia et al., 2021). Regarding bacterial communities, a combination of 4 forward and 2 reverse primers in equal concentration (**Table S5**) was used, targeting the V4 region of the 16S rRNA. For fungal communities, 6 forward primers and one reverse primer in equal concentration were used, targeting the ITS2 rRNA region (**Table S5**). To avoid the amplification of plant material, a mixture of peptide nucleic acid (PNA) probes inhibiting the plant mitochondrial (mPNA) and chloroplast DNA (pPNA) for 16S libraries, and a third mix of PNA blocking the plant ITS rRNA (itsPNA) (Lundberg et al., 2013) were used. Regarding AMF, a two-step PCR procedure to amplify the large ribosomal subunit (LSU) DNA was used, following to the protocol of Brígido et al., (2017). The specific primers LR1 and NDL22 (Van Tuinen et al., 1998) were used in the first PCR, whereas the primers FRL3 and FRL4 (Gollotte et al., 2004) were used to amplify the LSU-D2 rRNA genes of AMF in the second PCR (**Table S5**). All primers used to generate the microbial libraries (16S, ITS and 28S) contained an extension used in PCR2 for the tagging with specific sequences to allow the future identification of each sample. As well, PCR-s were prepared without addition of fungal DNA (negative control) and on known fungal and bacterial communities (mock communities) as quality controls. The amplicons were visualised by electrophoresis through a 1% agarose gel in 1X TBE buffer. PCR products were purified using the Agencourt AMPure XP PCR purification kit (Beckman Coulter), following the manufacturer protocol. After DNA purification, PCR products were quantified with a Qubit^®^2.0 fluorometer (Invitrogen) and new PCRs performed for samples with concentration lower than 2.5 ng.µL^-1^. Samples with DNA concentration higher than 2.5 ng.µl^-1^ were sent for tagging (PCR2) and MiSeq Illumina next-generation sequencing (GenoScreen for ITS and 28S, PGTB INRAE for 16S).

### Sequence processing

After sequences demultiplexing and barcode removal, fungal, bacterial and glomerales sequences were processed using FROGS (Find Rapidly OTU with Galaxy Solution) (Escudié et al., 2018) implemented on the Galaxy analysis platform (Afgan et al., 2016). Sequences were clustered into OTUs based on the iterative Swarm algorithm, and then chimaeras and fungal phiX contaminants were removed. As suggested by Escudié and collaborators (Escudié et al., 2018), OTUs with a number of reads lower than 5.10^-5^ percent of total abundance, and not present in at least 3 samples, were removed. Fungal sequences not assigned to the ITS region using the ITSx filter implemented in FROGS were then discarded and fungal sequences were affiliated using the UNITE Fungal Database (Nilsson et al., 2019), the bacterial sequences using SILVA database and 28S glomerales sequences using MaarjAM database (Öpik et al., 2010). OTUs with a BLAST identity lower than 90% and BLAST coverage lower than 95% were considered as chimaeras and removed from the dataset. Additionally, sequences affiliated with chloroplasts and mitochondria were removed. In order to achieve an equal number of reads in all samples, the rarefy_even_depth function from Phyloseq package (McMurdie and Holmes, 2013) in R (R Core Team. 2020) was used. To optimize the analyses of fungal community structures and diversity, a different rarefaction threshold was applied depending on microbial communities. We rarefied bacterial communities with a number of sequences to 4377, 5139 for fungal communities (ITS) and 6831 for 28S communities. FUNGuild (Nguyen et al., 2016) and FungalTraits (Põlme et al., 2020) were combined to classify each fungal OTU into an ecological trophic guild. A confidence threshold was applied to only keep “highly probable” and “probable” affiliated trophic guilds and the other OTUs were assigned as “unidentified”.

### Metabolite profiling

Untargeted metabolite levels were determined from lyophilized roots and shoots as described in Tschaplinski et al., (2012). To ensure complete extraction, freeze-dried, powdered material (∼25 mg for shoot samples and 30 mg for root samples) was twice extracted overnight with 2.5 mL of 80% ethanol (aqueous), sorbitol (75 µL (L) or 50 µL (R and Myc) of a 1 mg/mL aqueous solution) was added to the first extract as an internal standard to correct for subsequent differences in derivatization efficiency and changes in sample volume during heating. The extracts were combined, and 500-µL (L) or 2-mL (R and Myc) aliquots were dried under nitrogen. Metabolites were silylated to produce trimethylsilyl derivatives by adding 500 µL of silylation-grade acetonitrile to the dried extracts followed by 500 µL of N-methyl-N-trimethylsilyltrifluoroacetamide with 1% trimethylchlorosilane and heating for 1 h at 70°C. For lyophilised root exudates, sorbitol (10 µL; 1 mg*ml-1) was added as internal standard prior to drying under nitrogen and silylating as described above but using 200 µL of each silylation solvent and reagent. After 2 days, a 1-µL aliquot was injected into an Agilent Technologies (Santa Clara, CA) 7890A/5975C inert XL gas chromatograph / mass spectrometer (MS) configured as previously described (Tschaplinski et al., 2012). The MS was operated in electron impact (70 eV) ionization mode using a scan range of 50-650 Da. Metabolite peaks were quantified by area integration by extracting a characteristic mass-to-charge (m/z) fragment with peaks scaled back to the total ion chromatogram using predetermined scaling factors and normalized to the extracted mass, the recovered internal standard, the analyzed volume and the injection volume. The peaks were identified using a large in-house user-defined database of ∼2700 metabolite signatures of trimethylsilyl-derivatized metabolites and the Wiley Registry 12^th^ Edition combined with NIST 2020 mass spectral database. The combination of these databases allowed accurate identification of a large fraction of the observed metabolites. Unknowns were designated by their retention time (min) and key m/z. The assignation of the distinct metabolic pathways was performed using the Kyoto Encyclopedia of Genes and Genomes database (KEGG) conjointly with the Plant Metabolic Network (PMN) focusing on *Populus trichocarpa* (https://pmn.plantcyc.org/POPLAR).

### Statistical analyses

To perform statistical analyses and data representation, the R software (R Core Team. 2016) was used. Soil parameters were tested for normal distribution using Shapiro Wilk tests. If the data were normally distributed, the differences between the means were assessed using Student t-tests followed by the Bonferroni correction, otherwise Wilcoxon tests were used. The difference of root and shoot fresh weight between poplar grown on natural and sterilised soil was assessed using a Wilcoxon test followed by Bonferroni corrections. The dynamics of root exudation over time was assessed using a Kruskal-Wallis test with false discovery rate (FDR) corrections. The differences among sampling time were assessed with a Fisher LSD post-hoc test. The differences of metabolites between natural and gamma-irradiated soil among plant organs were assessed using a Wilcoxon test followed by FDR corrections. Finally, multiple regression using redundancy analysis (RDA, *rda* function in vegan package) were used for bacterial and fungal communities between root and shoot habitats and in the rhizosphere over time with plant metabolites and root exudates as explanatory variables. The significance of plant metabolites and root exudates and their correlations with microbial communities were assessed using the *envfit* function in vegan with 1000 permutations and applied FDR corrections. An ANOVA-like permutation test (function *anova.cca* in the vegan package with 1,000 permutations) was then used to determine if RDA models were statistically significant. Differences in fungal and bacterial community structures between tissues and time were tested using permutational multivariate analysis of variance (PERMANOVA, *adonis2* function in vegan package) based on Bray-Curtis and Jaccard distances, and differences in structures were visualised using a nonmetric dimensional scaling (NMDS) ordination. The significance of microbial communities and environmental variables and their correlations were calculated using the *envfit* function in vegan with 1,000 permutations and applied FDR corrections. The difference of richness and diversity between genotypes over time was assessed using the Kruskal-Wallis test, with Bonferroni corrections followed by the Fisher LSD post-hoc test. The difference of fungal and bacteria relative abundance at the Phylum and genera level between organs was tested using Kruskal-Wallis tests, with Bonferroni corrections for Phylum and FDR correction for genera. In order to reduce the weight of the correction on fungal genera, only fungal genera with a relative abundance higher than 1% were kept and a Kruskal-Wallis test was applied, followed by Bonferroni correction. The variations of fungal trophic guilds relative abundance were assessed by Kruskal-Wallis tests, followed by Bonferroni correction, while fungal diversity and richness were analysed using Kruskal-Wallis tests and LSD post-hoc tests.

### Data availability

Raw data were deposited in the NCBI Sequence Read Archive (SRA) under SRA accession numbers SRR26346063 to SRR26346115 for the 16S data, SRR26286627 to SRR26286662 to for ITS data and SRR26286064 to SRR26286106 for 28S data (project PRJNA1017804).

## Acknowledgements

FF was supported by “Contrat Doctoral” from the Lorraine Université d’Excellence. This research was sponsored by the Plant-Microbe Interfaces Scientific Focus Area in the Genomic Science Program, the Office of Biological and Environmental Research in the U.S. Department of Energy, Office of Science. Oak Ridge National Laboratory is managed by UT-Battelle, LLC, for the U.S. Department of Energy, Office of Science (DE-AC05-00OR22725), LABEX ARBRE (ANR-11-LABX-0002-01).

## Author Contributions

FF, CVF and AD designed and coordinated the research and the experimental design. Poplar *in vitro* cultures were produced and maintained by FF, FG. Sampling was realised by FF and FG. DNA extractions were done by FF. Root exudates and plant metabolome characterization were performed by NLE and TJT. Data analyses were performed by FF, TJT, CVF and AD. FF, AD and CVF wrote the manuscript. TJT helped revise the manuscript. All authors approved the final version of the manuscript.

