## Supplemental Figures for "Microbe tree metabolite interactions in the soil - phyllosphere continuum of poplar tree: when microbes rewire poplar root exudate and metabolome"

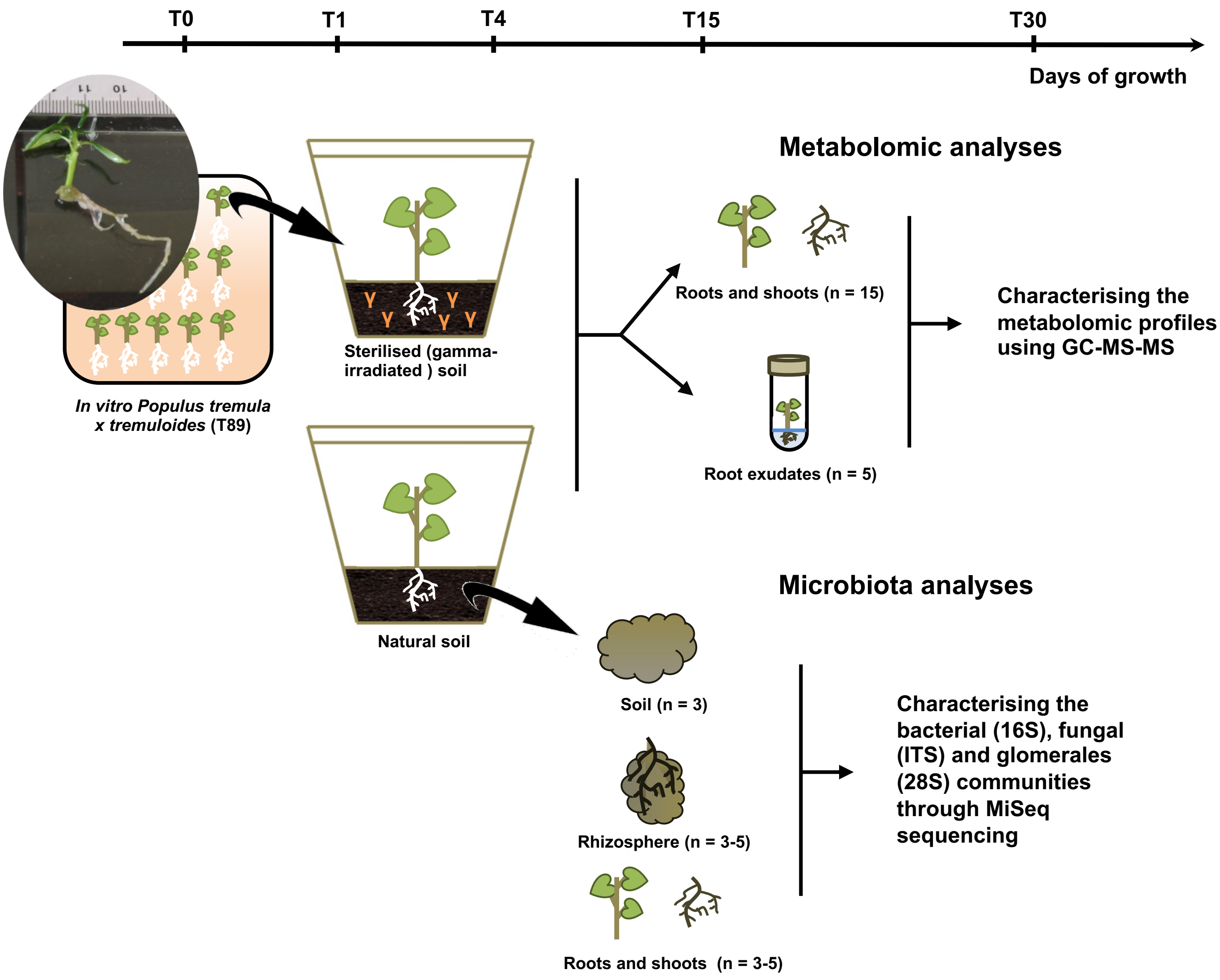


**Figure S1. Experimental design of the study.** Poplar cuttings (*Populus tremula x tremuloides* T89) were grown *in vitro* for 3 weeks before being transferred in microcosms containing either natural or sterilised (gamma-irradiated) soil and grown for 30 days. Before transplantation, roots and shoots (n = 3) were sampled to confirm the axenic status of cuttings. After 1, 4, 15 and 30 days, we sampled the soil (n = 3), the rhizosphere (n = 3-5), the roots (n = 3-5) and shoots (n = 3-5) of poplar grown in natural soil for microbial communities analyses. In parallel, we collected the root exudates (n = 5) after 4, 15 and 30 days of growth as well as the roots (n = 15-25) and shoots (n = 15-25) after 30 days of poplar grown in natural and sterilised soil for metabolomic analyses. At each sampling time, we measured the fresh biomass for both roots and shoots.

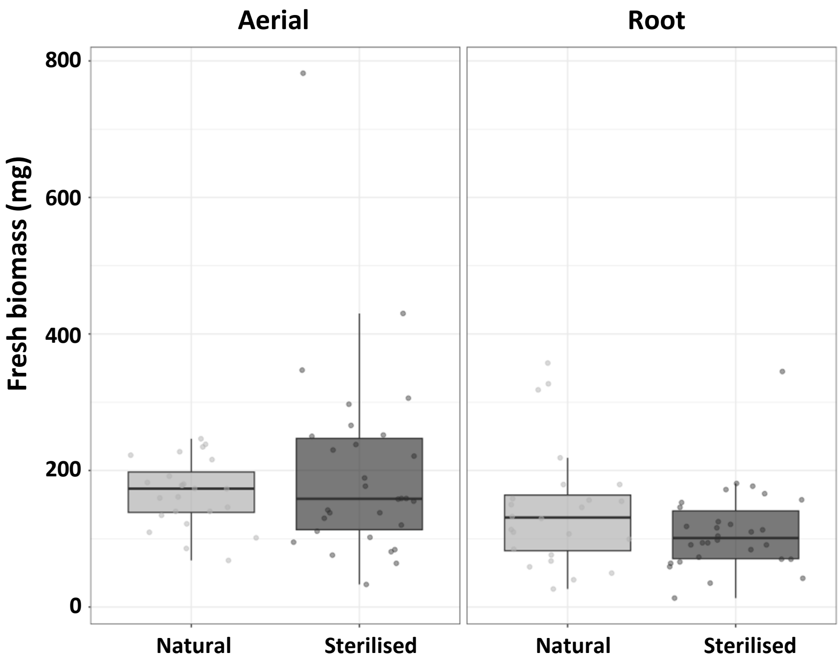


**Figure S2. Influence of microorganisms on poplar growth after 30 days.** No significant difference of aerial and root growth of poplar grown in natural and sterilised soil over 30 days (n = 15-25, Wilcoxon, Bonferroni corrected, p.adj > 0.05).

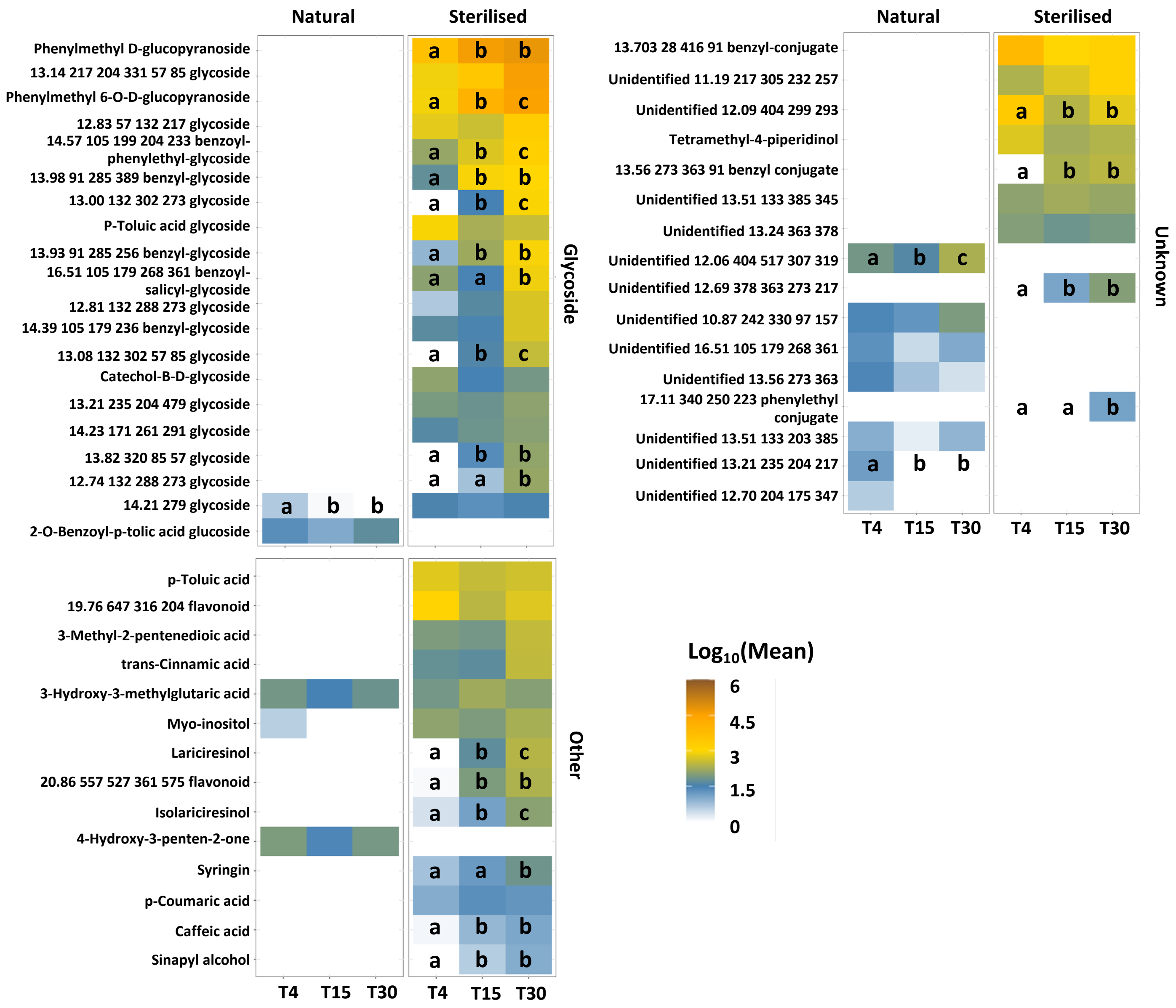


**Figure S3. Influence of microorganisms on the root exudates profile over time.** Dynamics of root exudates of poplar grown in presence or absence of microorganisms. Values correspond to the exudate mean concentration transformed by Log10. Letters indicate significant differences of metabolite concentration over time for each treatment (n = 5, Kruskal-Wallis, FDR corrections, p.adj ≤ 0.05, Fisher LSD post-hoc test).

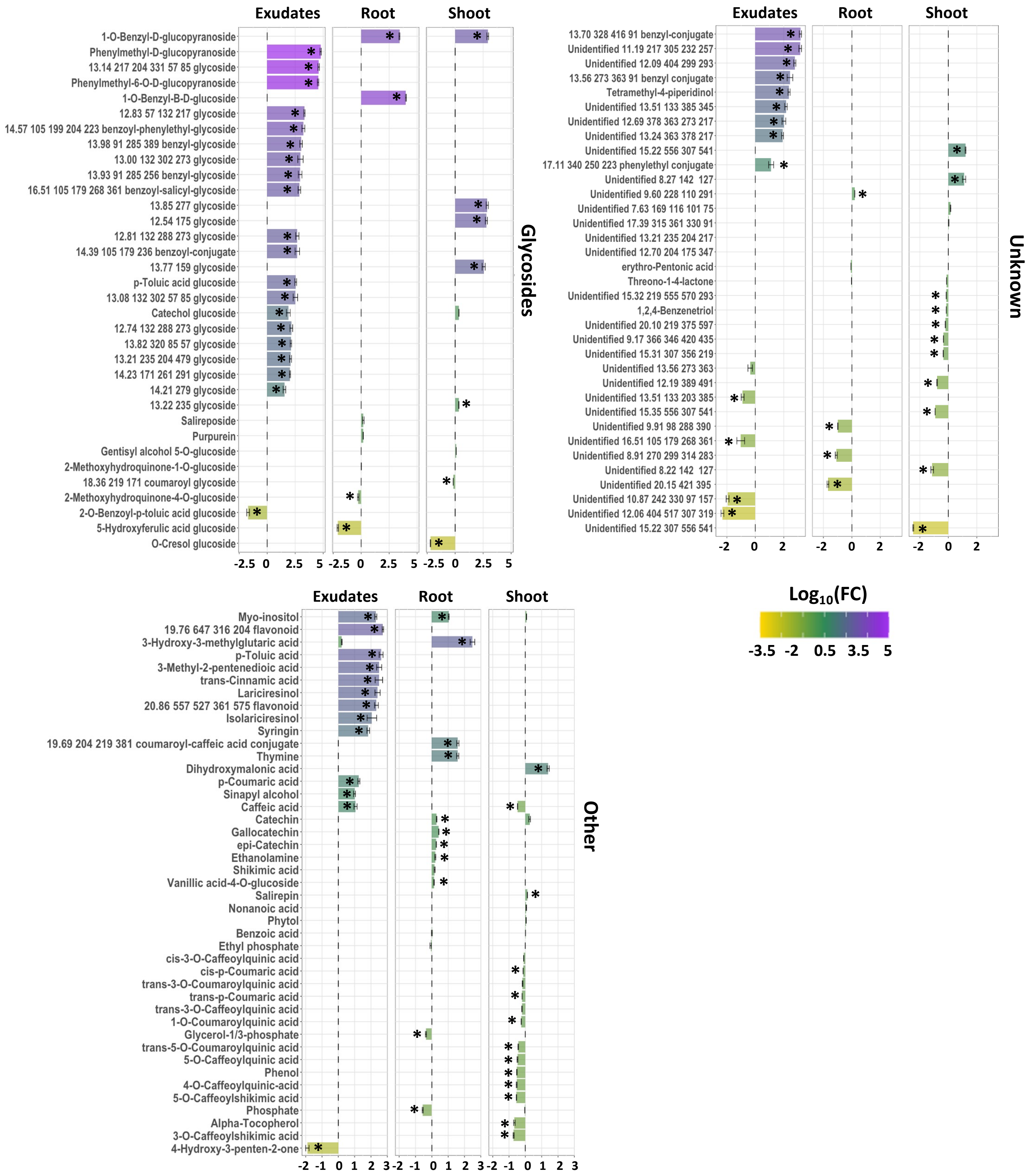


**Figure S4. Influence of microorganisms on the composition and abundance of root exudates, root and shoot metabolites after 30 days of growth.** Bars represent the log2 fold change of the relative abundance of metabolic compounds detected in sterilised soil (positive bars) versus natural soil (negative bars). * indicate significant difference of metabolite abundance between the two treatments (n = 5-25, Wilcoxon, FDR corrections, p.adj ≤ 0.05).

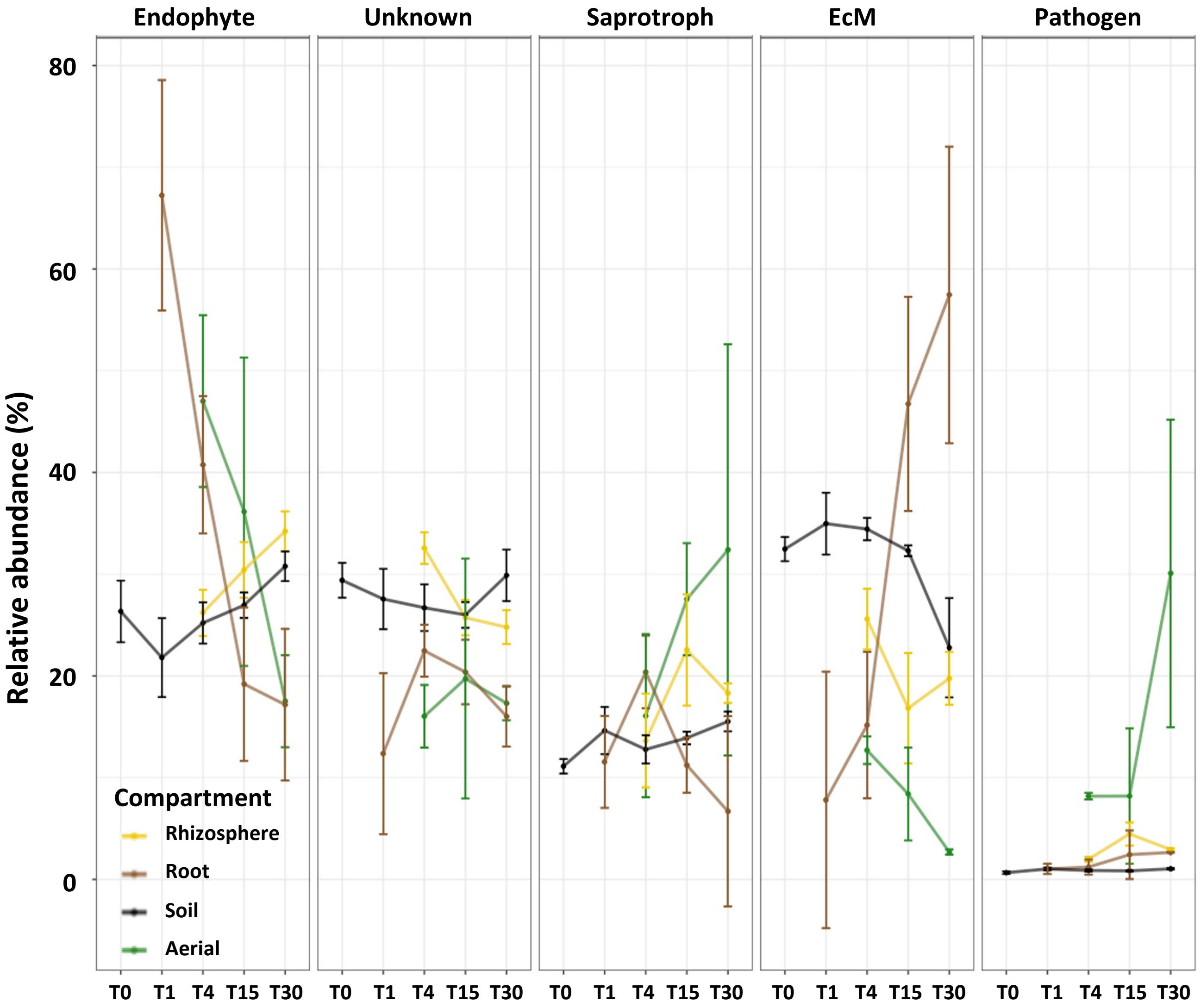


**Figure S5. Relative abundance of the different fungal trophic guilds assigned between the soil, rhizosphere, root and shoot over 30 days of growth**. Ecological trophic guilds assignation was performed using the FUNGuild (Nguyen et al., 2016) and FungalTraits (Põlme et al., 2020) databases. Kruskal-Wallis, Bonferroni correction, p.adj > 0.05, n = 3-5. Points represent the mean relative abundance of each guild and bars indicate their standard error.

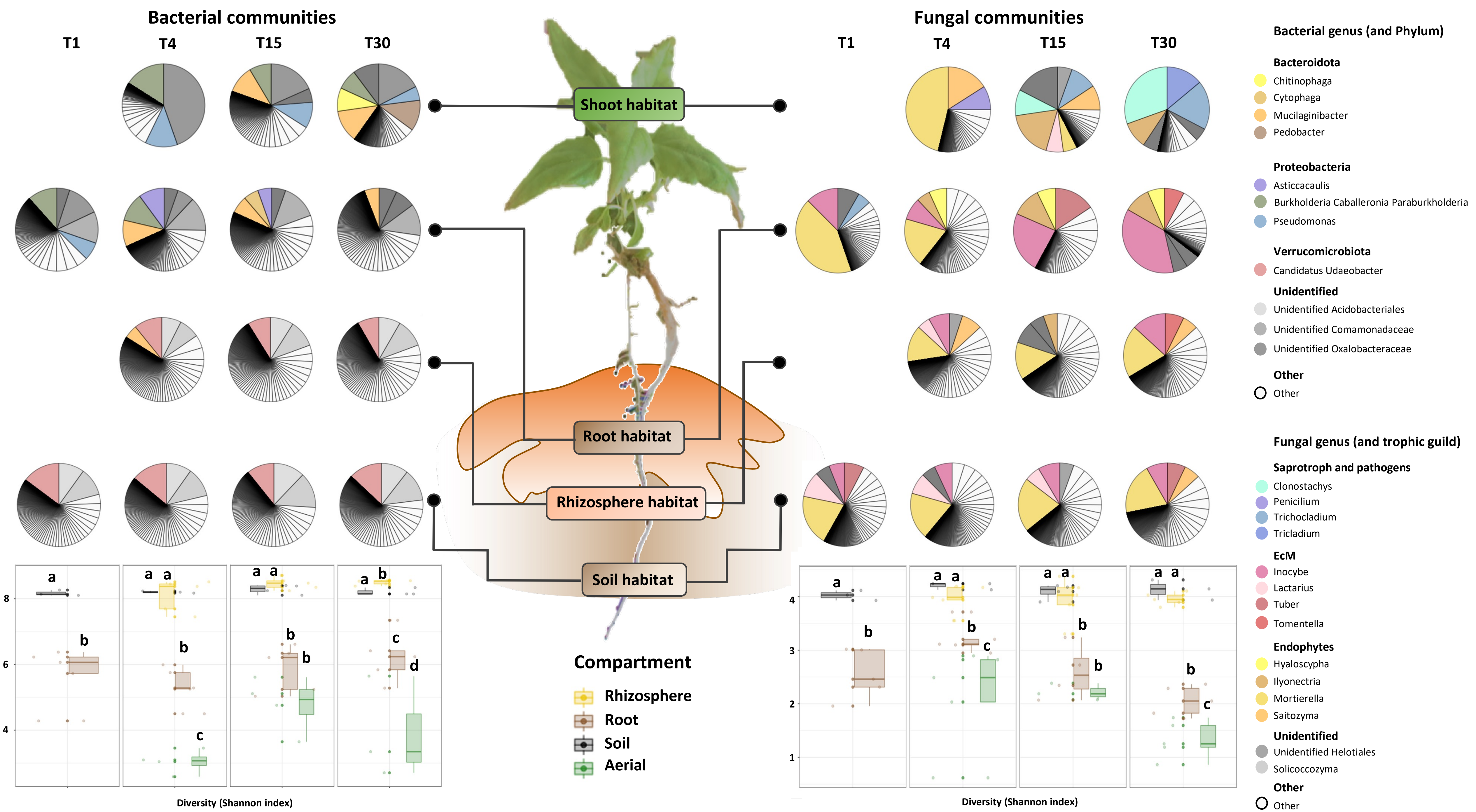

Bacterial communities
T1
T4
T15
T30
Fungal communities
T1
T4
T15
T30
Shoot habitat
Root habitat
Rhizosphere habitat
Soil habitat
Compartment
Rhizosphere
Root
Soil
Aerial
Diversity (Shannon index)
Diversity (Shannon index)
Bacterial genus (and Phylum)
Bacteroidota
Chitinophaga
Cytophaga
Mucilaginibacter
Pedobacter
Proteobacteria
Asticcacaulis
Burkholderia Caballeronia Paraburkholderia
Pseudomonas
Verrucomicrobiota
Candidatus Udaeobacter
Unidentified
Unidentified Acidobacteriales
Unidentified Comamonadaceae
Unidentified Oxalobacteraceae
Other
Other
Fungal genus (and trophic guild)
Saprotroph and pathogens
Clonostachys
Penicilium
Trichocladium
Tricladium
EcM
Inocybe
Lactarius
Tuber
Tomentella
Endophytes
Hyaloscypha
Ilyonectria
Mortierella
Saitozyma
Unidentified
Unidentified Helotiales
Solicoccozyma
Other
Other

**Figure S6. Relative abundance of the dominant (>1%) and diversity (Shannon index) of bacterial and fungal communities across habitat over 30 days of growth.** Relative abundance of (A) bacterial and (B) fungal genera and their diversity in the four compartments sampled (soil, rhizosphere, root and shoot), (Kruskal-Wallis, FDR correction, p.adj $\leq$ 0.05, n = 3-5).

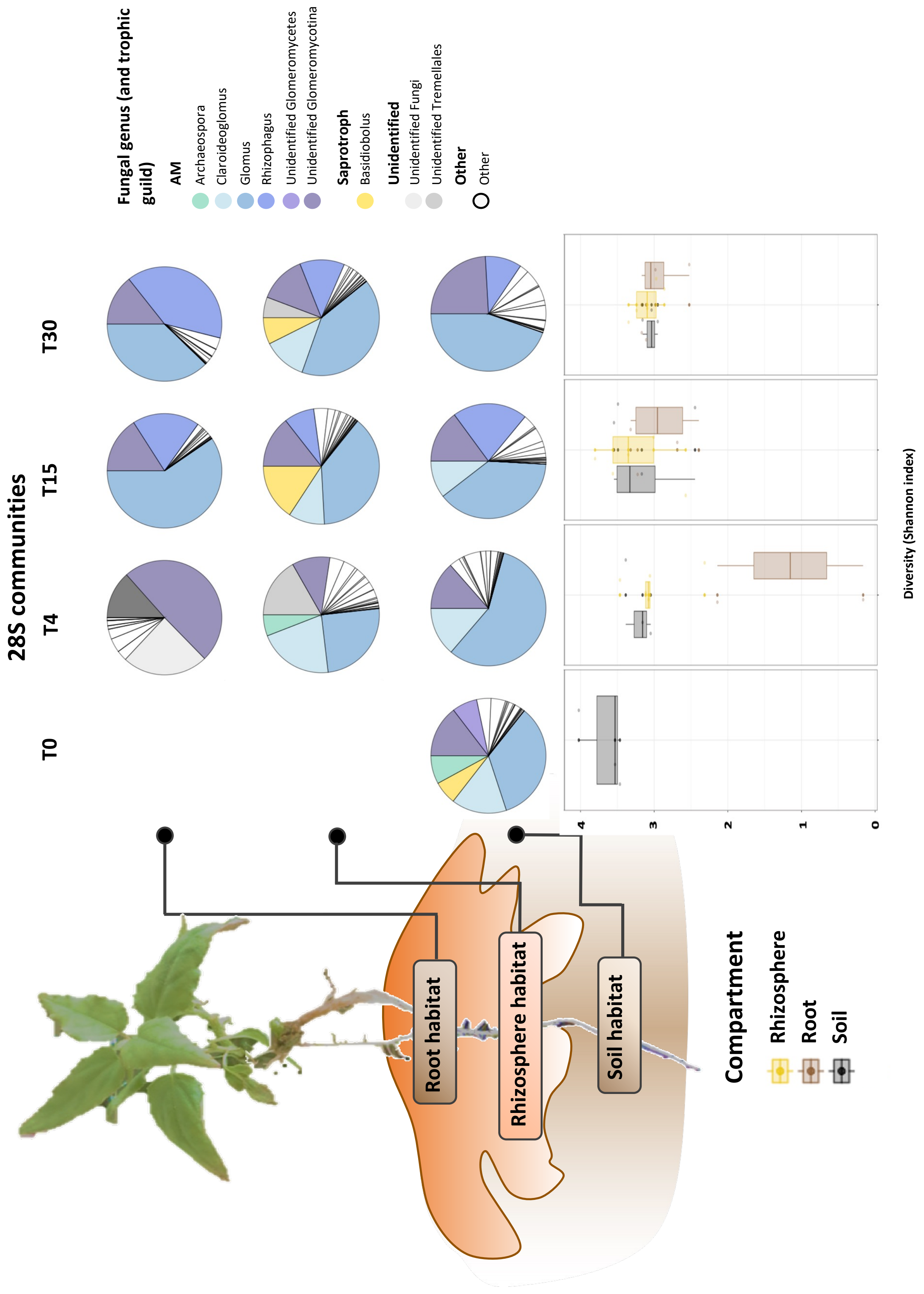


**Figure S7. Relative abundance of the dominant (>1%) and diversity (Shannon index) of 28S communities across habitat over 30 days of growth.** Glomerales genera and their diversity in the 3 compartments sampled (soil, rhizosphere, and root), (Kruskal-Wallis, FDR correction, p.adj ≤ 0.05, n = 3-5).

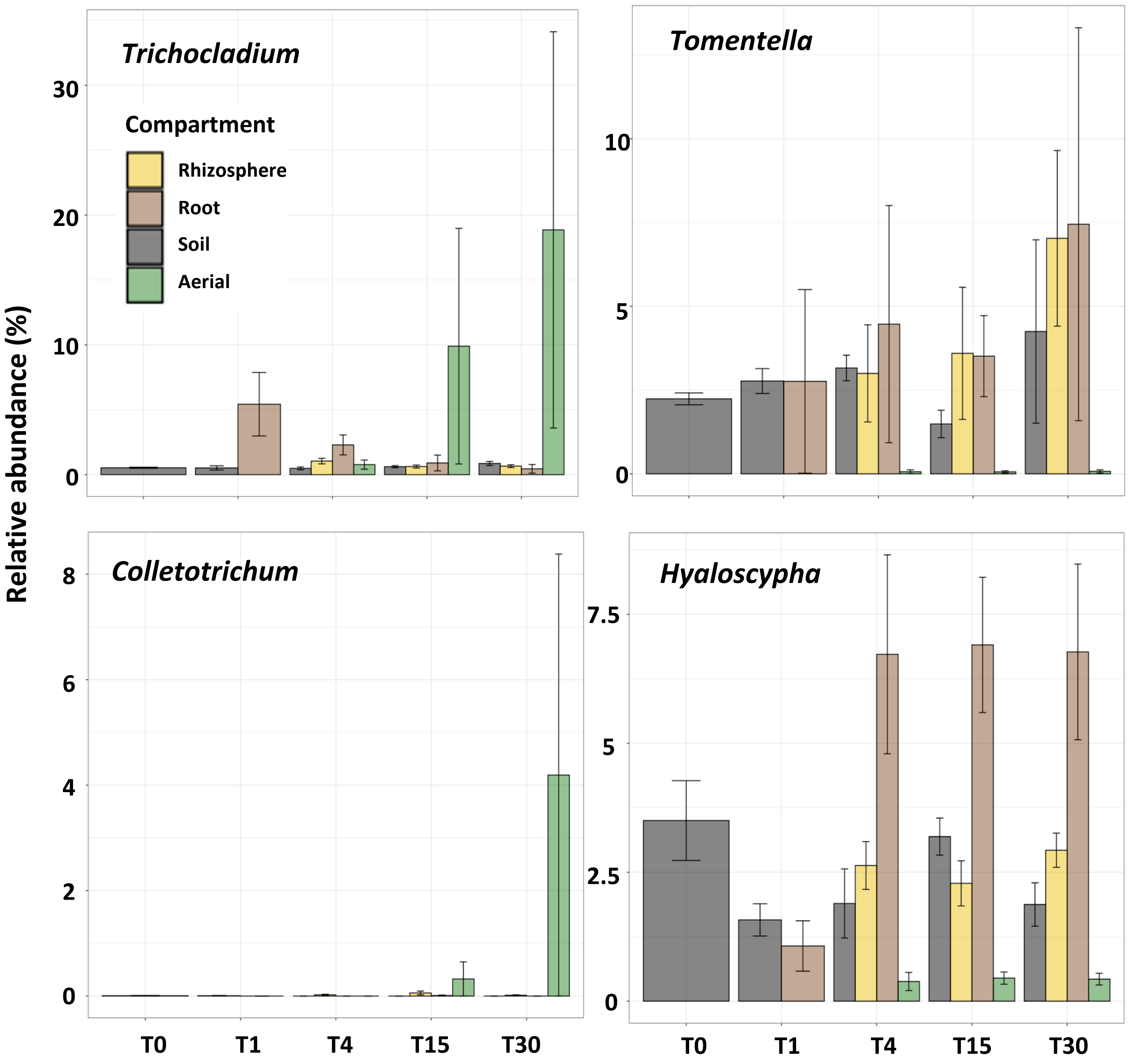


**Figure S8. Relative abundance of fungal genera associated with specific habitat over 30 days of growth.** Fungal taxa were chosen according to their significance related to particular time or habitat in multivariate partition analyses after multiple regression analyses and 1,000 permutation (FDR corrected, p.adj ≤ 0.01). Histograms represent the mean relative abundance of each taxa and bars indicate their standard error (n = 3-5).

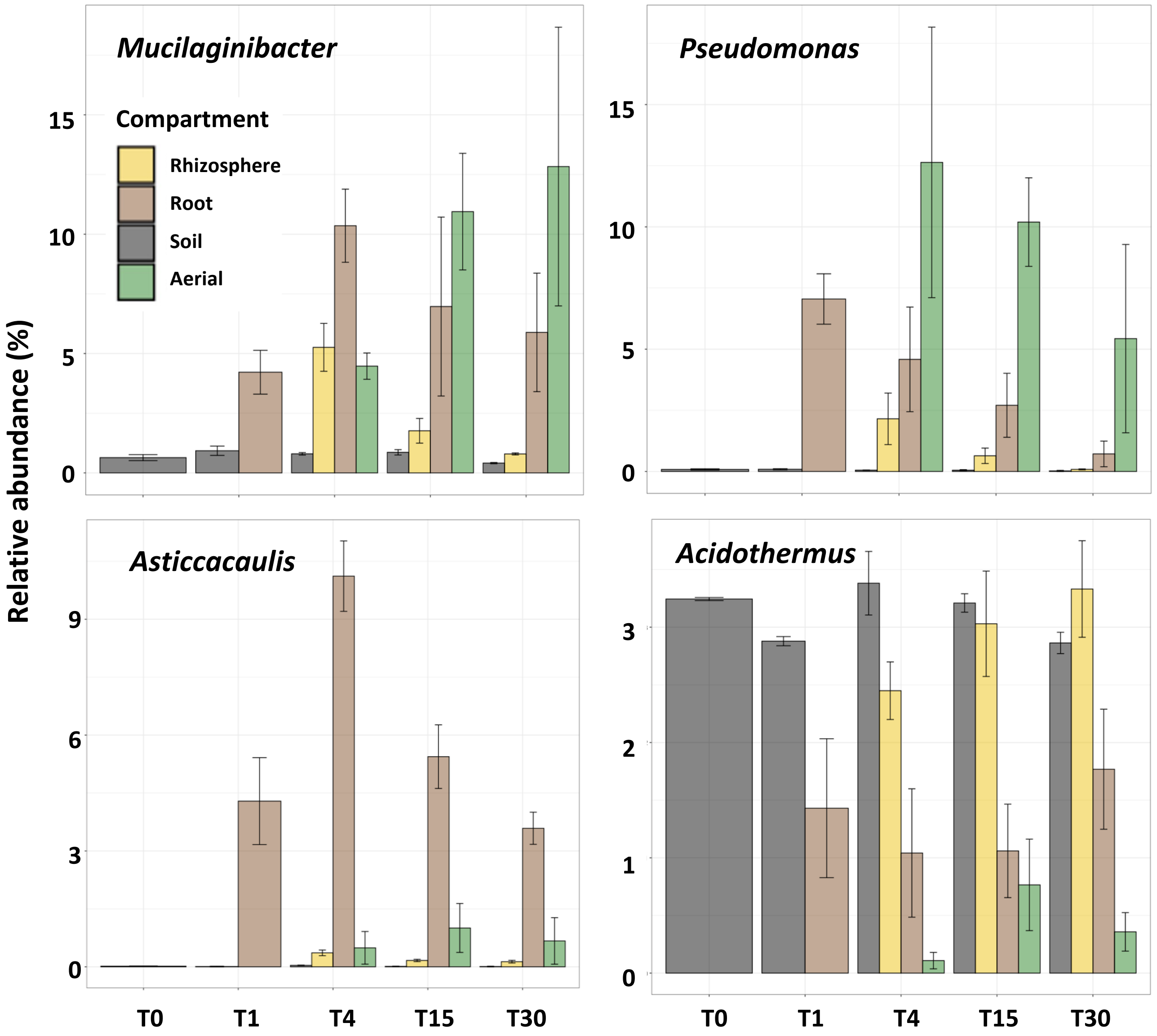


**Figure S9. Relative abundance of bacterial genera associated with specific habitat over 30 days of growth.** Bacterial taxa were chosen according to their significance related to particular time or habitat in multivariate partition analyses after multiple regression analyses and 1,000 permutation (FDR corrected, p.adj ≤ 0.01). Histograms represent the mean relative abundance of each taxa and bars indicate their standard error (n = 3-5).
